# Genotyping-by-sequencing-based identification of *Arabidopsis* pattern recognition receptor RLP32 recognizing proteobacterial translation initiation factor IF1

**DOI:** 10.1101/2021.03.04.433884

**Authors:** Li Fan, Katja Fröhlich, Eric Melzer, Isabell Albert, Rory N. Pruitt, Lisha Zhang, Markus Albert, Sang-Tae Kim, Eunyoung Chae, Detlef Weigel, Andrea A. Gust, Thorsten Nürnberger

## Abstract

Pattern-triggered immunity (PTI) is a central component of plant immunity. Activation of PTI relies on the recognition of microbe-derived structures, termed patterns, through plant encoded surface-resident pattern recognition receptors (PRRs). We have identified proteobacterial translation initiation factor 1 (IF1) as an immunogenic pattern that triggers PTI in *Arabidopsis thaliana* and some related Brassicaceae species. Unlike most other immunogenic patterns identified, IF1 elicitor activity cannot be assigned to a small peptide epitope, suggesting that tertiary fold features are required for IF1 receptor activation. We have deployed natural variation in IF1 sensitivity to identify leucine-rich repeat (LRR) receptor-like protein 32 (RLP32) as the corresponding *Arabidopsis* receptor using a restriction site-associated DNA sequencing (RAD-seq) approach. Transgenic expression of RLP32 confers IF1 sensitivity to *rlp32* mutants, IF1-insensitive *Arabidopsis* accessions and IF1-insensitive *Nicotiana benthamiana*. RLP32 binds IF1 specifically and forms complexes with LRR receptor kinases SOBIR1 and BAK1 to mediate signaling. Similar to previously identified PRRs RLP32 confers resistance to *Pseudomonas syringae* infection, highlighting an unexpectedly complex array of bacterial pattern sensors within a single plant species.

---

Metazoans and plants employ innate immune systems to cope with microbial infections. Immunogenic microbe-derived signatures from pathogenic, commensal or beneficial microbes, collectively referred to as microbe or pathogen-associated molecular patterns (MAMPs/PAMPs), serve as ligands for host-encoded cell surface pattern recognition receptors (PRRs)^1,2^. Pattern recognition and subsequent initiation of intracellular immune signaling culminates in the activation of antimicrobial defenses ultimately restricting pathogen spread.

In plants, pattern-triggered immunity (PTI) controls attempted infections by host non-adapted microbes and contributes to basal immunity against host-adapted pathogens^1–4^. PTI suppression by pathogen-derived effectors is an element of successful infection of host plants by host-adapted microbes. Effector-mediated host susceptibility has driven the evolution of intracellular immune receptors that recognize effector activities on host plant targets and mediate activation of immunity to host-adapted pathogens, a process termed effector-triggered immunity (ETI)^5,6^. Mutual potentiation of PTI and ETI pathways in *Arabidopsis thaliana* (hereafter *Arabidopsis*) has been proposed, suggesting mechanistic links between these two layers of plant immunity^7,8^. An *Arabidopsis* plasma membrane-associated intracellular signaling complex linking helper NLRs (NUCLEOTIDE-BINDING LEUCINE-RICH REPEAT RECEPTORS) from the ADR1 (ACTIVATED DISEASE RESISTANCE 1) family and the lipase-like proteins EDS1 (ENHANCED DISEASE SUSCEPTIBILITY 1) and PAD4 (PHYTOALEXIN DEFICIENT 4) to plant PRRs may provide a convergence point for PTI and ETI signaling^9^.

Plant cell surface-resident PRRs are distinguished by structurally diverse extracellular domains for ligand binding, including leucine-rich repeat (LRR), lysin-motif (LysM) or lectin domains^1,2^. LRR-domain proteins predominantly mediate the perception of microbe-derived proteins or peptides^10^, and are classified as either LRR receptor kinases (LRR-RKs) or LRR receptor proteins (LRR-RPs) depending on the presence or absence of a cytoplasmic kinase domain^1,2^. LRR-RPs form constitutive heteromeric complexes with the adaptor kinase SUPPRESSOR OF BRASSINOSTEROID INSENSITIVE 1 (BRI1)-ASSOCIATED KINASE (BAK1)-INTERACTING RECEPTOR KINASE 1 (SOBIR1) and, like LRR-RKs, bind to members of the SOMATIC EMBRYOGENESIS RECEPTOR KINASE (SERK) protein family in a ligand-dependent fashion^11^. PRR complex formation subsequently triggers downstream signaling pathways which, though overlapping, differ depending on the receptor^12^.

In the past two decades, several plant PRRs recognizing molecularly defined patterns from bacteria, fungi and oomycetes have been identified^1,2,4^. In addition, immune-stimulating insect or parasitic plant-derived patterns and their cognate immune sensors have been elucidated^13–15^. *Arabidopsis* LRR-RK FLAGELLIN SENSING 2 (FLS2) recognizes fragments of bacterial flagellins containing a 22-amino-acid motif (flg22)^16^. FLS2 activities have been found throughout higher plants, which is in contrast to the majority of PRRs exhibiting plant genus or species-specific distribution^1^. Other *Arabidopsis* LRR-type sensors for bacteria-derived patterns include ELONGATION FACTOR THERMO-UNSTABLE RECEPTOR (EFR), XANTHINE/PERMEASE SENSING 1 (XPS1) and RLP23^17–19^. As for FLS2, small immunogenic epitopes within these patterns have been defined, including elf18 (from ELONGATION FACTOR THERMO-UNSTABLE), xup25 (from XANTHINE PERMEASE) and nlp20 (from NECROSIS AND ETHYLENE-INDUCING PROTEIN 1-LIKE PROTEINs)^16,18,19^. Bacteria-derived pattern sensors have also been found in other plant species. These include *Nicotiana benthamiana* and tomato receptors CSPR (CSP22 RESPONSIVENESS) and CORE (COLD SHOCK PROTEIN RECEPTOR) for bacterial cold shock protein fragment csp22 and tomato FLS3, which recognizes a flagellin fragment (flgII-28) unrelated to flg22^20–22^.

Accumulating evidence suggests that plants employ multiple PRRs to sense a given microbe^1^. For example, *Arabidopsis* harbors FLS2, EFR and XPS1 to recognize three *Pseudomonas syringae*-derived peptide patterns^16,18,19^ in addition to two immune sensors recognizing bacterial medium-chain 3-hydroxy fatty acids and peptidoglycans^23,24^. This complexity is likely to increase given that the *Arabidopsis* genome encodes more than 600 transmembrane receptor-like proteins^25–27^. Beside an evident academic interest that drives the identification of plant PRRs and their microbe-derived patterns, such receptors may also be used to engineer durable disease resistance in crop plants. Interfamily transfer of plant PRRs into crops has been demonstrated to confer novel pattern recognition capabilities and enhanced immunity to infection by host-adapted pathogens^28^. Thus, PRR combinations employed in transgenic crops may become an important tool to reduce crop losses and to secure global food security.

Here, we report the identification of *Arabidopsis* RLP32 as a sensor of proteobacterial protein translation initiation factor 1 (IF1). Unlike for other PRRs, RLP32 activation requires the IF1 tertiary fold to mediate IF1 recognition and immunity to bacterial infection in *Arabidopsis* and in RLP32-transgenic *N. benthamiana*. Our findings indicate a striking diversity of bacterial pattern recognition systems in one particular plant.

## Results

### *Ralstonia solanacearum*-derived pattern recognition in *Arabidopsis*

Plant pathogenic *R. solanacearum* has previously been reported to produce *Arabidopsis* defense elicitors other than bacterial flagellin^29^. In agreement with this, we found that protein fractions from liquid culture-grown *R. solanacearum* elicited plant defenses in the *Arabidopsis fls2 efr* mutant (Supplementary Fig. 1), suggesting that *R. solanacearum* elicitor activity (RsE) is not only different from flagellin-derived flg22, but also from elf18. RsE induces the production of reactive oxygen species (ROS), callose and the plant hormone ethylene, the phosphorylation of mitogen-activated protein kinases MPK3 and MPK6, as well as enhanced expression of the defense marker gene, *PATHOGENESIS-RELATED 1* (*PR1*) in *Arabidopsis* leaves (Supplementary Fig. 1). Proteinase K treatment abolished RsE immunogenic activity, suggesting that the elicitor is a peptide or protein (Supplementary Fig. 2a). In gel filtration experiments, the molecular mass of RsE was estimated to be < 10 kDa (Supplementary Fig. 2b).

To identify the RsE receptor, we screened a collection of 106 natural strains, or accessions, of *Arabidopsis* for RsE-induced ethylene production (Supplementary Fig. 3a). Three accessions (Dog-4, ICE21, and ICE73) with reproducibly strongly reduced ethylene production relative to that of reference accession Col-0 were deemed RsE-insensitive and selected for in-depth analysis (Fig. 1a, Supplementary Fig. 3a). These accessions remained sensitive to SCLEROTINIA CULTURE FILTRATE ELICITOR 1 (SCFE1)^30^, an unrelated fungal elicitor recognized by RLP30 (Fig. 1a), suggesting that insensitivity to RsE was not due to a general defect in defense activation. Approximately half of the tested accessions produced more ethylene than Col-0 in response to RsE, including hypersensitive accession ICE153 (Fig. 1a, Supplementary Fig. 3a).

**Figure 1.**
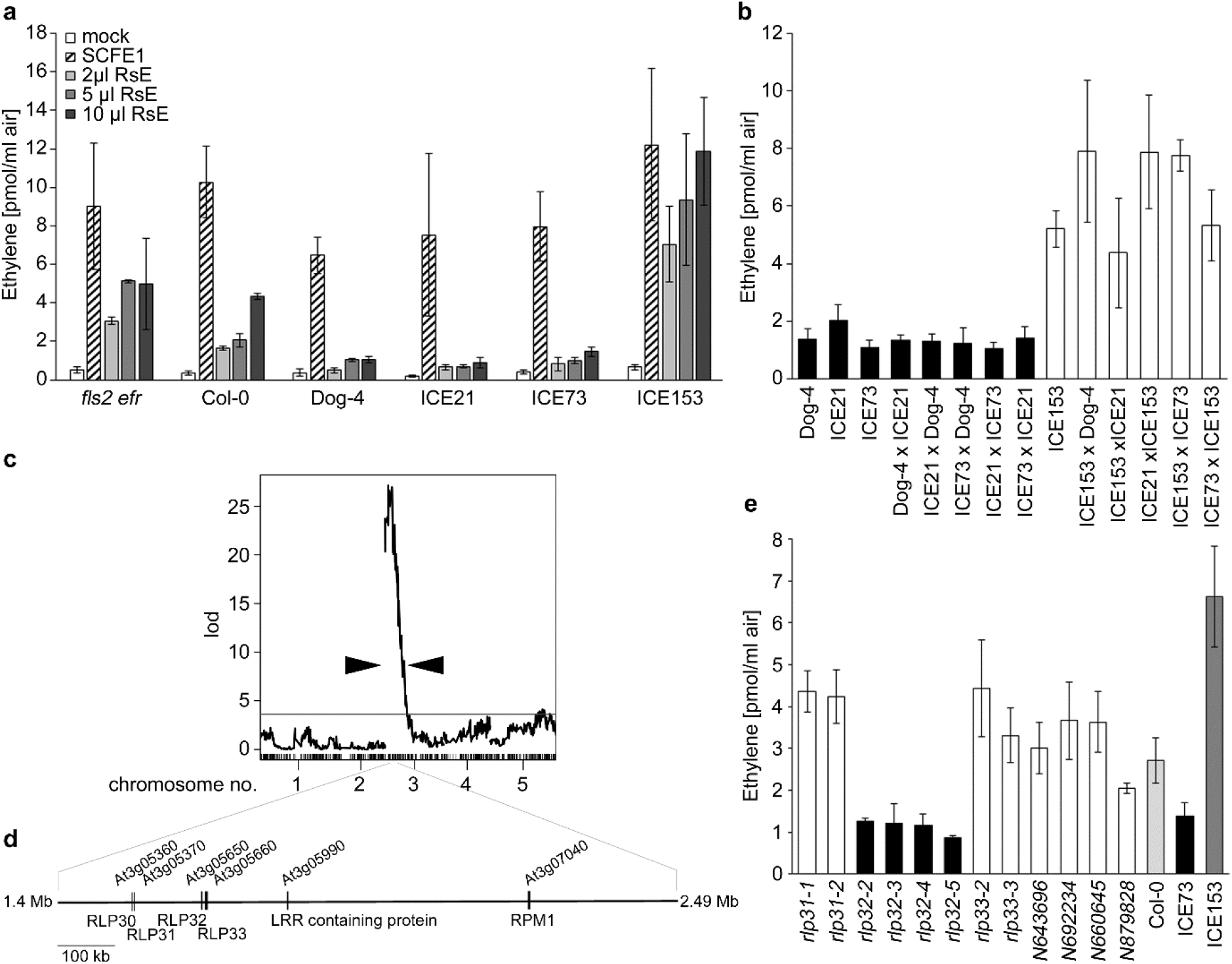
Rad-seq-associated QTL analysis-based identification of the RLP32 locus conferring sensitivity to *R. solanacearum* elicitor RsE. **a,** Ethylene production in *Arabidopsis* Col-0 wild-type, *fls2 efr* mutant and RsE-insensitive (Dog-4, ICE21, ICE73) and RsE-hypersensitive (ICE153) accessions. Treatments with water (mock) and SCFE1 served as controls. Shown are mean values of two replicates ± SD. **b**, RsE-induced ethylene production in crosses of sensitive and insensitive accessions as described in Supplementary Fig. 4. Black bars indicate insensitive accessions or a cross of two insensitive accessions, and white bars indicate sensitive accession ICE153 or its crosses with insensitive accessions. Shown are mean values of three replicates ± SD. **c,** rQTL mapping for RsE-induced ethylene response in F2 mapping populations of an ICE153 × ICE73 cross. LOD scores from a full genome scan across five chromosomes of *Arabidopsis* using a binary trait model for RsE-elicited ethylene scores. The grey line indicates a genome-wise α equaling to 0.05 LOD thresholds, which defines significant QTLs based on 1,000 permutations. Arrow heads indicate a region with LOD value above 10. **d,** Genomic arrangement of putative loci conferring RsE sensitivity within a 1.1 Mb region on chromosome 3 identified by QTL analysis. **e,** RsE-induced ethylene production in *Arabidopsis* plants carrying T-DNA insertions in genes indicated in (d). Ethylene production in Col-0 wild-type and accessions ICE73 and ICE153 served as controls. Shown are mean values of three replicates ± SD.

Insensitive accessions were crossed reciprocally (Fig. 1b, Supplementary Fig. 4). F_1_ progeny from all crosses remained RsE-insensitive, suggesting that changes at the same locus render these accessions insensitive to RsE. Crossing insensitive accessions with hypersensitive ICE153 produced only RsE-sensitive F_1_ plants (Fig. 1b). F_2_ populations of an ICE153 × ICE73 cross exhibited a segregation ratio of 1:3 (92 insensitive plants vs. 303 sensitive plants), suggesting that RsE sensitivity segregates as a single, recessive Mendelian trait. Thus, natural variation in *Arabidopsis* RsE sensitivity could be employed to identify the corresponding PRR.

### Genotyping-by-sequencing-based identification of RLP32

Restriction site-associated DNA sequencing (RAD-seq) was conducted to identify the genomic region conferring RsE sensitivity in *Arabidopsis*. DNA samples of 84 RsE-insensitive and 108 RsE-sensitive F_2_ plants of the ICE153 × ICE73 cross were digested with *Pst*I/*Mse*I, and a tagged DNA fragment library was single-end sequenced (Illumina HiSeq2000, 36,4-fold average coverage) to yield 16,973 potential markers. 901 markers were selected for further quantitative trait locus (QTL) mapping. Whether using quantitative or binary traits (RsE-sensitive 0, RsE-insensitive 1), QTL analyses identified one major peak on chromosome 3 associated with RsE sensitivity, consistent with Mendelian segregation (Fig. 1c, Supplementary Fig. 5a). Confidence interval analysis (LOD score 10) using a binary trait model defined a QTL interval between markers Chr3:20373 and Chr3:6582498 (Supplementary Fig. 5b). Recombination breakpoint analyses in F_2_ populations derived from the ICE153 × ICE73 cross were used to narrow down the QTL to a 1.1 Mb region flanked by markers Chr3:1399533 and Chr3:2485009, respectively (Supplementary Fig. 6). This region contained 339 open reading frames, 6 of which encoded LRR-type proteins as prime candidates for PRRs (RLP30, At3g05360; RLP31, At3g05370; RLP32, At3g05650; RLP33, At3g05660; LRR-containing protein At3g05990; disease-related R protein/RPM1, At3g07040) (Fig. 1d). We focused on the analysis of LRR proteins first as proteinaceous immunogenic patterns, such as RsE (Supplementary Fig. 2a), are predominantly recognized by LRR ectodomain receptors. RLP30 was excluded from further analysis because RsE-insensitive accessions recognized SCFE1 (Fig. 1a) and because SCFE1-insensitive accession Bak-2 was responsive to RsE (Supplementary Fig. 3b). Knockdown or knockout alleles of the remaining genes were tested for RsE-inducible ethylene production. Four independent T-DNA/transposon alleles of RLP32 (insertions validated by flanking fragment sequencing; Supplementary Fig. 7), proved insensitive to RsE, suggesting that RLP32 confers RsE sensitivity (Fig. 1e). All other tested mutants responded to RsE (Fig. 1e).

Col-0 RLP32 is composed of an ectodomain comprising 23 LRR units with an island domain separating LRRs 19 and 20, a juxta-membrane domain, a trans-membrane domain and a 28-amino-acid-tail (Supplementary Fig. 8). Inspection of RLP32 protein sequences in RsE-insensitive accessions revealed rather diverse polymorphisms within RLP32 orthologs (Supplementary Fig. 9), thus making predictions about mutations causal for loss of RsE-sensitivity difficult. Since RsE-insensitive accessions are distantly related (Supplementary Fig. 10) and because RsE-insensitive RLP32 proteins are distinguished by substantial sequence polymorphisms, we conclude that loss of RsE pattern recognition may have occurred several times independently.

### RLP32 recognizes proteobacterial translation initiation factor 1 (IF1)

To assess whether RsE activity is found in bacteria other than *R. solanacearum*, protein extracts from plant non-pathogenic *Escherichia coli* were fractionated using the protocol for RsE preparation. As shown in Supplementary Fig. 11, Col-0 wild-type plants, but not *rlp32* mutants produced ethylene upon *E. coli* elicitor treatment, suggesting that RsE activity is not restricted to *R. solanacearum*.

Partially purified *E. coli* elicitor was subjected to liquid chromatography mass spectrometry (LC-MS/MS)-based protein identification (Supplementary Fig. 11). Of a total of 290 proteins identified, 20 proteins were chosen for further analysis according to the following criteria: (i) a high number of peptides with different masses found in all four elicitor-active fractions, (ii) a predicted molecular mass of RsE of approximately 10 kDa (Supplementary Fig. 2b), and (iii) a basic isoelectric point as deduced from the migration of RsE elicitor activity in ion exchange chromatography experiments. Of the candidate proteins produced in *E. coli*, only protein translation initiation factor 1 (IF1) induced RLP32-dependent ethylene production (Fig. 2a, Supplementary Fig. 11c).

**Figure 2.**
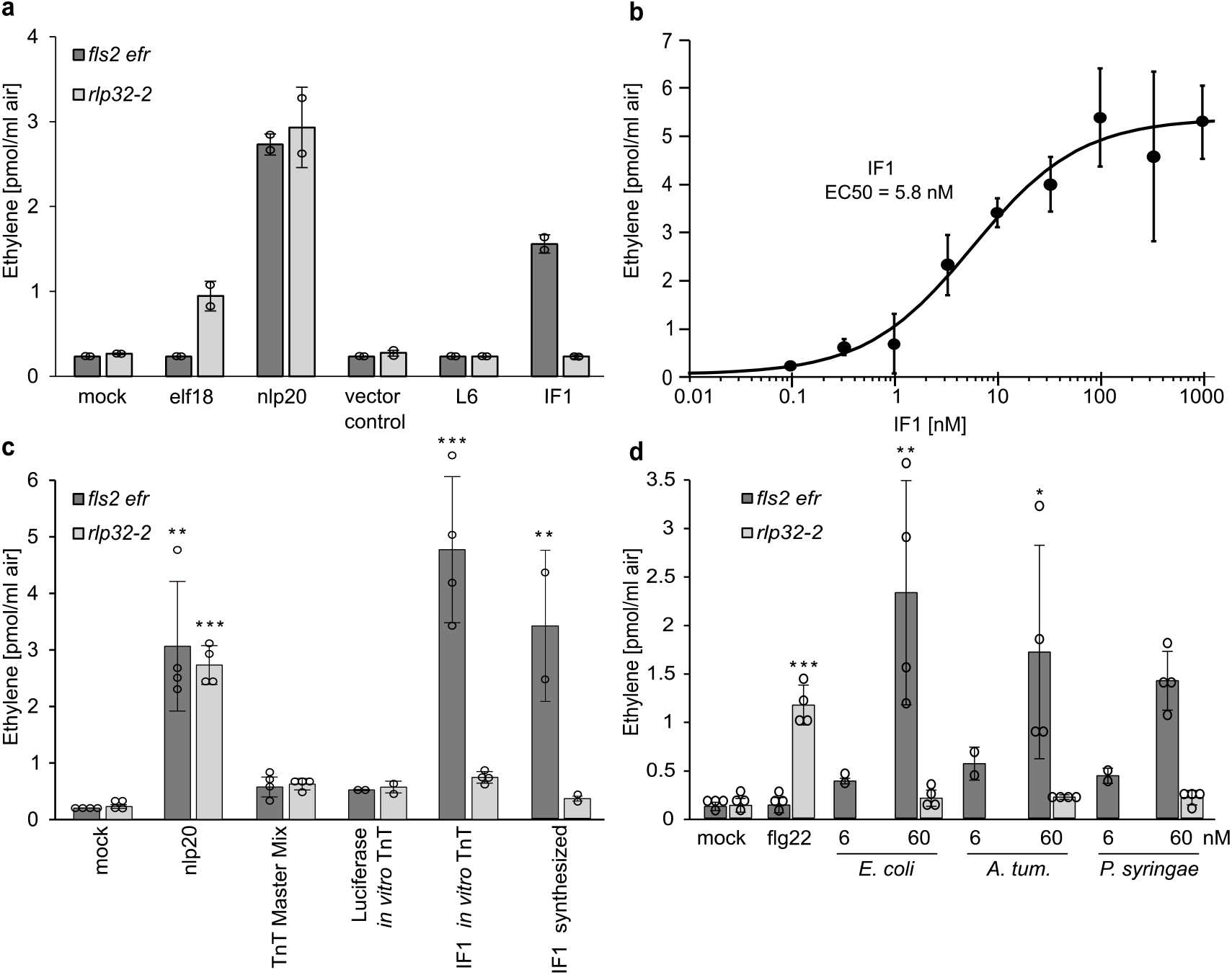
IF1 elicitor activity. **a**, Ethylene accumulation was determined in *Arabidopsis fls2 efr* or *rlp32* mutants treated with recombinant IF1 and ribosomal protein L6, or with proteins prepared from *E. coli* transformed with empty vector. Treatment with water (mock), elf18 or nlp20 served as controls. **b**, Determination of EC_50_ values using increasing concentrations of recombinant IF1 (produced in yeast). EC_50_ values and curve fit were calculated using 4P Rodbard Model comparison. **c**,**d**, Ethylene accumulation in *Arabidopsis fls2 efr or rlp32* mutants treated with *in vitro* translated (TnT) IF1, chemically synthesized IF1 (**c**), or recombinant IF1 from the bacterial species indicated. *A. tum*, *Agrobacterium tumefaciens*. (**d**). Bars represent means ± SD of two (**a,c,d**), three (**b**) or four (**c**, TnT-IF and **d**, each pooled from two experiments) replicates. (* *P*≤0,5, ** *P*≤0,01, *** *P*≤0,001, Dunnetts’s test with mock (**a, d**) or TnT MasterMix (**c**) as control, respectively). Experiments were performed at least twice, with similar results.

IF1 is a 71-amino-acid single domain protein composed of a five-stranded β-barrel and a short α-helical loop separating strands 3 and 4 (Supplementary Fig. 12). IF1 belongs to the oligonucleotide-binding-fold protein family and shares close structural homology to another member of this family, bacterial cold shock protein CspA^31^, a trigger of immunity in tobacco and tomato^32^. Recombinant *E. coli* IF1 triggers ethylene production at low nanomolar concentrations (EC_50_ = 5.8 nM) (Fig. 2b). Likewise, *E. coli* IF1 produced by *in vitro* transcription and translation or by chemical synthesis exhibited RLP32-dependent elicitor activity (Fig. 2c). This finding indicates that IF1, and not a copurifying contaminant, triggers RLP32-mediated plant defense.

IF1 amino acid sequences are highly conserved amongst proteobacteria (Supplementary Fig. 12a). IF1 proteins from plant pathogenic *Pseudomonas syringae*, *Agrobacterium tumefaciens* and *R. solanacearum* share with *E. coli* IF1 85 %, 60% or 75% protein sequence identity, respectively. Likewise, I-TASSER-based 3-dimensional structure prediction revealed strong secondary and tertiary structure conservation of plant pathogenic bacteria-derived IF1 molecules when compared to *E. coli* IF1 (Supplementary Fig. 12b). IF1 preparations from these bacterial species exhibited elicitor activities similar to that of *E. coli* IF1 (Fig. 2d, Supplementary Fig. 13). Our findings suggest that IF1 is a widespread bacterial pattern and that *R. solanacearum* IF1 accounts for the elicitor activity in RsE preparations (Supplementary Fig. 13).

Stable expression of *pRLP32::RLP32* constructs in *Arabidopsis rlp32* mutants and in the RsE-insensitive accession ICE73 conferred sensitivity to both RsE and IF1 (Fig. 3a,b). Likewise, over-expression of *p35S::RLP32-GFP* in RsE- and IF1-insensitive *N. benthamiana* conferred sensitivity to IF1 (Fig. 3c). Collectively, these data confirm that the elicitor RsE is IF1 and that RLP32 mediates IF1-inducible defenses without the need for additional species-specific factors.

**Figure 3.**
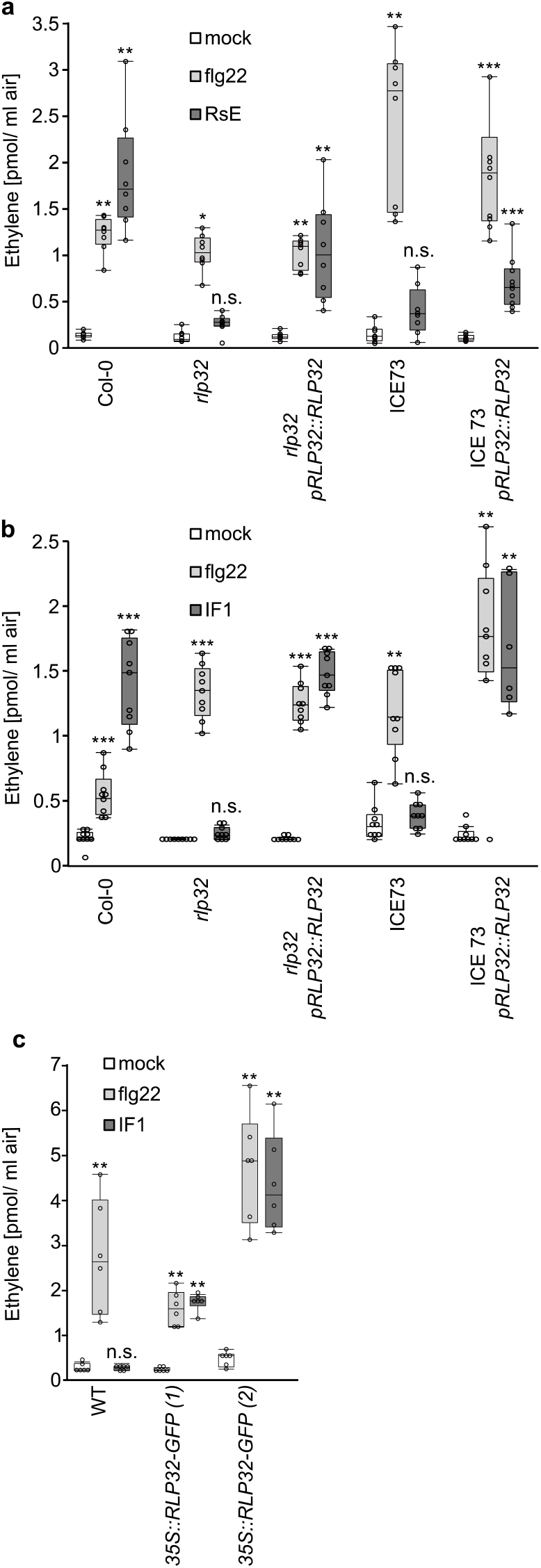
RLP32 confers sensitivity to RsE and IF1. **a - d,** Ethylene accumulation after treatment with flg22, IF1 (expressed with 6xHis in *E. coli*), RsE or water (mock) in *Arabidopsis rlp32* mutants, RsE-insensitive accession ICE73, *Arabidopsis* plants stably expressing *pRLP32::RLP32* (**a,b**), and in two independent *p35S::RLP32–GFP* transgenic *N. benthamiana* lines (**c**). Bars represent means ± SD (**a**, *n*=8; **b**, *n*=9; **c**, *n*=6, pooled from each two to three experimental repetitions; * *P*≤0,5, ** *P*≤0,01, *** *P*≤0,001, n.s. not significant, Steel test with mock treatment as control).

### Tertiary structure properties are required for IF1-mediated plant defense activation

Immunogenic activities of large proteinaceous patterns are typically represented by short, conserved peptide fragments. We have tested chemically synthesized nested peptides spanning the entire IF1 protein sequence, as well as recombinantly expressed IF1 fragments carrying N-terminal or C-terminal deletions (Fig 4a,b). Peptide fragments were designed to preserve IF1 secondary structure motifs (Supplementary Fig. 12d). With the exception of a near full-length IF1 variant (I6-R71) with a N-terminal deletion of a short unstructured segment of five amino acid residues (Supplementary Fig. 12b), all IF1 peptide fragments failed to trigger RLP32-dependent ethylene production (Fig. 4a,b). These findings suggest that tertiary structure rather than primary or secondary structure motifs determine IF1 elicitor activity.

**Figure 4.**
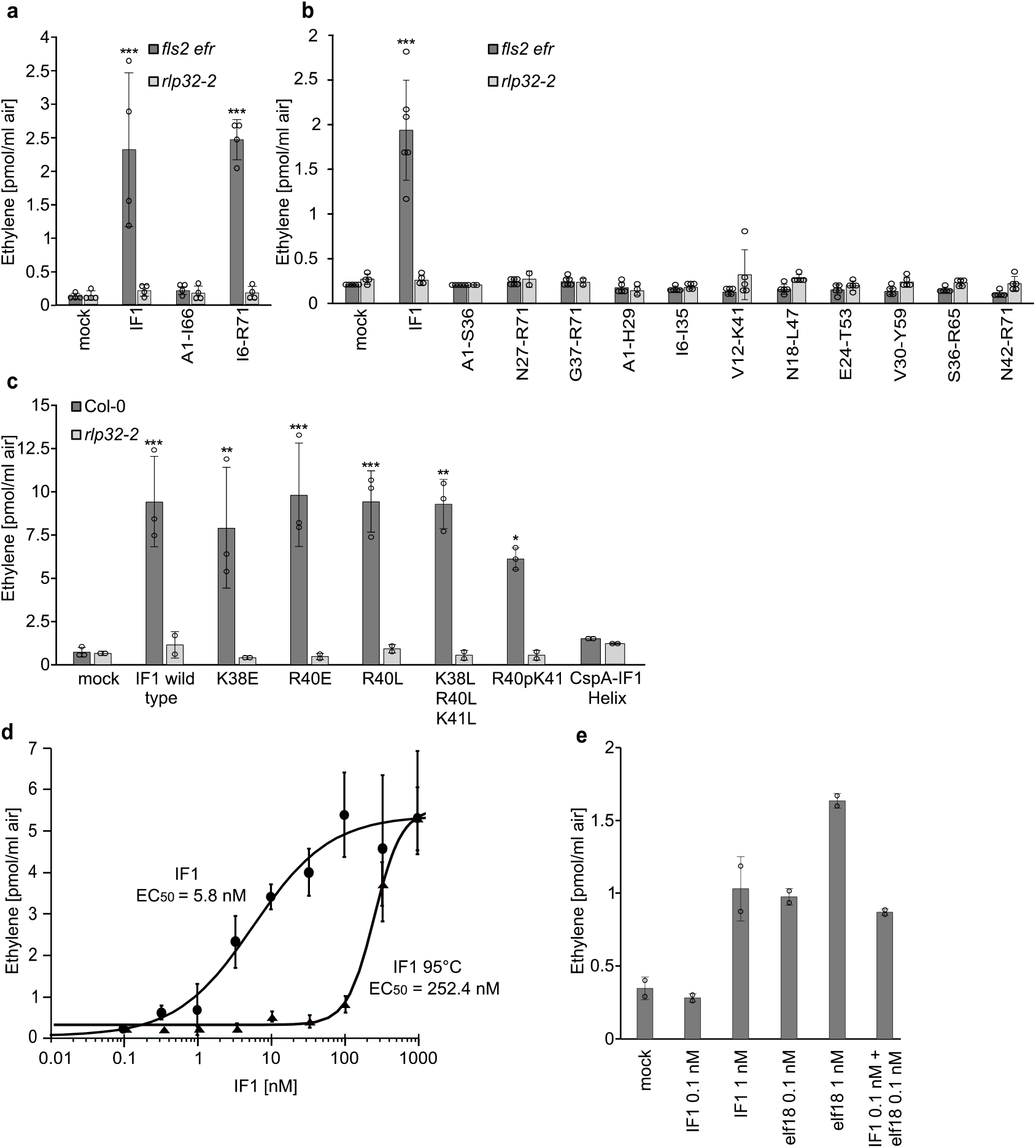
IF1 tertiary fold features are required for its elicitor activity. **a**, Ethylene accumulation in *Arabidopsis fls2 efr or rlp32-2* mutants treated with 60 nM IF1 or with 40 nM IF1_I6-R71 and IF1_A1-I66 (produced in *E.coli)*, respectively. **b**, Ethylene accumulation after treatment with IF1 or indicated synthetic IF1 variants. **c**, Ethylene accumulation after treatment with IF1 produced in yeast, indicated IF1 point mutants or a CspA-IF1 Helix chimeric protein. **d**, Determination of EC_50_ values using increasing concentrations of IF1 produced in yeast before and after heat treatment. **e**, Ethylene accumulation in *Arabidopsis* Col-0 wild-type plants treated with IF1 and elf18 alone or in combinations indicated. Bars represent means ± SD (**a**, *n*=4; **b**, *n*≥4; **c**, *n*≥2; **d**, *n*=3, **a,b**, pooled data from each two experiments; * *P*≤0,5, ** *P*≤0,01, *** *P*≤0,001, n.s. not significant, Dunnett’s test with mock treatment as control)

Proteobacterial IF1 and cold shock protein CspA share a highly conserved five-stranded β-barrel fold^31^. One notable difference is that the α-helical motif separating IF1 β-strands 3 and 4 is absent in CspA. As *E. coli* CspA is not recognized by *Arabidopsis* (Supplementary Fig. 14), we hypothesized that this short helix is important for IF1 elicitor activity. Simultaneous replacement of positively charged residues K38, R40 and K41 by leucine residues or introduction of a proline residue, an amino acid known to distort helical structures because of forming a kink in the peptide backbone, did not reduce IF1 elicitor 7 activity (Fig 4c). Likewise, introduction of the IF1 helical motif into the 5-stranded β-barrel of *E. coli* CspA did not restore elicitor activity in the chimeric protein (Fig. 4c). Hence, the IF1 helical motif is likely not important for IF1 elicitor activity. Heat treatment strongly reduced IF1 elicitor activity (Fig. 4d), suggesting that heat-induced alterations within the IF1 tertiary fold adversely affected its ability to trigger plant defense. Altogether, our findings suggest that tertiary structure features rather than primary or secondary structure motifs determine IF1 elicitor activity.

### IF1 and elf18 do not act additively

IF1 and EF-Tu are distinct components of the proteobacterial protein translation machinery that are perceived by *Arabidopsis* PRRs RLP32 and EFR, respectively. We have tested whether simultaneous action of both patterns, a scenario likely occurring during bacterial infection, has additive or synergistic effects on plant defense activation. IF1 and EF-Tu fragment elf18 were applied either alone or in combination at concentrations below or corresponding to the EC_50_ values of the respective patterns^18^ (Fig. 2b). However, neither additive nor synergistic effects on pattern-induced ethylene production were found (Fig. 4e).

### RLP32 is required for IF1-induced plant immunity

Pattern treatment primes plant immunity to subsequent infection by host-adapted, virulent plant pathogens. To test whether this also applies to IF1, we pre-treated Col-0 plants with IF1 24 hours before infection with virulent *Pseudomonas syringae* pv. *tomato* (*Pst*) strain DC3000 (Fig. 5a). We found that treatment with IF1, like the positive control nlp20, reduced *Pst*DC3000 growth 3 days post infection compared to mock-treated plants. This priming effect was abolished in two independent *rlp32* mutant lines (Fig. 5b,c), suggesting that IF1 recognition by RLP32 contributes to plant immune activation and reduced microbial proliferation on infected plants.

*N. benthamiana* plants are insensitive to IF1 treatment. However, stable transformation with *p35S::RLP32-GFP* conferred IF1 sensitivity to independent transgenic lines (Fig. 3d). When infected with bacterial strain *Pst* DC3000 hrcC^−^ transgenic plants exhibited reduced bacterial growth when compared to that observed in untransformed control plants (Fig. 5d).

**Figure 5.**
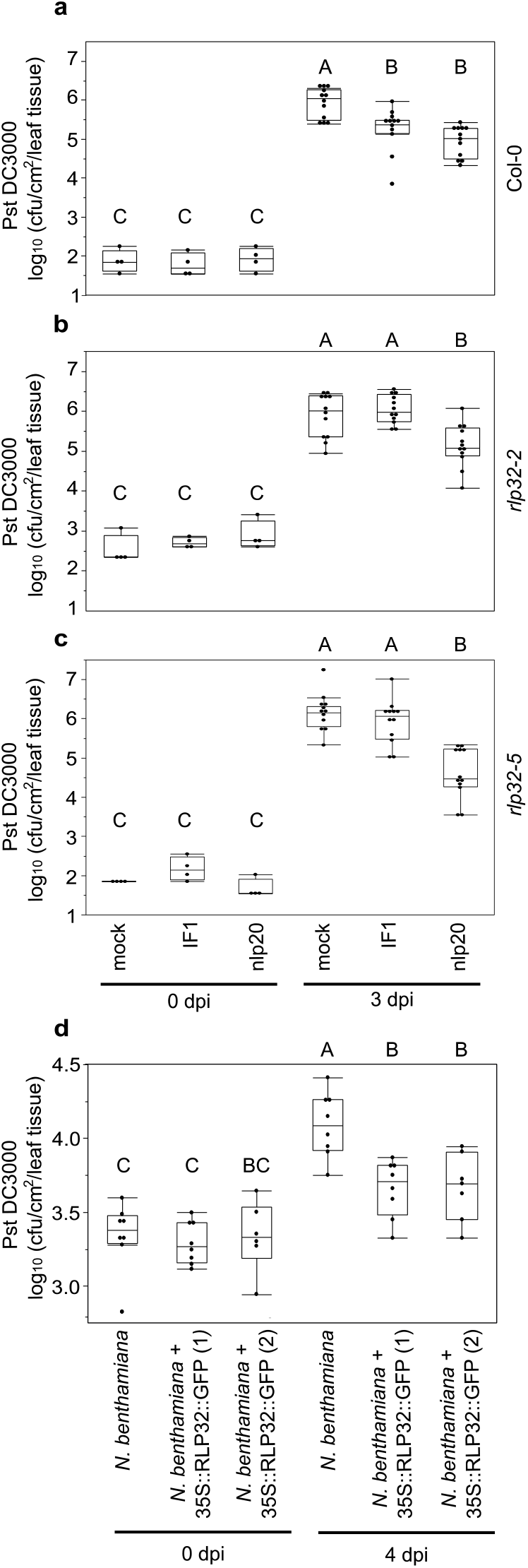
RLP32 mediates bacterial resistance. **a - c**, Growth of *Pseudomonas syringae* pv. *tomato* DC3000 (*Pst*) in *A. thaliana* Col-0 (**a**) and *rlp32* plants (**b,c**) after priming with IF1 or nlp20 24 h prior to bacterial infection. Water infiltration served as a control (mock). Bacterial growth was determined 0 and 3 days post infection (dpi). **d**, Growth of *Pst* DC3000 hrcC- in wild-type or two *RLP32*-transgenic *N. benthamiana* lines at 0 and 4 dpi. Box plots show the minimum, first quartile, median, third quartile, and a maximum of log cfu/cm^2^ (**a**, *n*=4 from 2 plants for 0 dpi and *n*=12 from 6 plants for 3 dpi; **b**, *n*=8 from 4 plants for 0 dpi and 4 dpi). Labels A-C indicate homogenous groups according to post-hoc comparisons fol**lowing multiple comparison analysis (a, b, c,** Steel-test with mock as a control; **d**, Steel-Dwass-test). Experiments were performed three times, with similar results.

### RLP32 binds IF1 and forms a PRR complex with co-receptors SOBIR1 and BAK1

Biologically active, biotinylated IF1 (bio-IF1) was employed to analyze binding to RLP32 *in planta* (Fig. 6, Supplementary Fig. 15). Leaves of *p35S::RLP32-GFP*-expressing *N. benthamiana* plants were treated with bio-IF1 before infiltration of the homo-bifunctional chemical cross-linker, EGS (ethylene glycol bis-succinimidyl succinate). RLP32-GFP was subsequently precipitated with GFP-trap beads and analyzed for ligand binding using a streptavidin-alkaline phosphatase conjugate. Control experiments were conducted using biotinylated nlp24 (nlp24-bio) cross-linked to transiently expressed RLP23-GFP^17^. In both cases, ligand binding to the respective receptors was observed at concentrations similar to those required for pattern-induced ethylene production (Fig. 6a). Loss of bio-IF binding to RLP32 in the presence of a 1,000-fold molar excess of native IF1 demonstrated ligand specificity of this binding event (Fig. 6a). Taken together, our findings suggest that RLP32 is a sensor for proteobacterial IF1.

**Figure 6.**
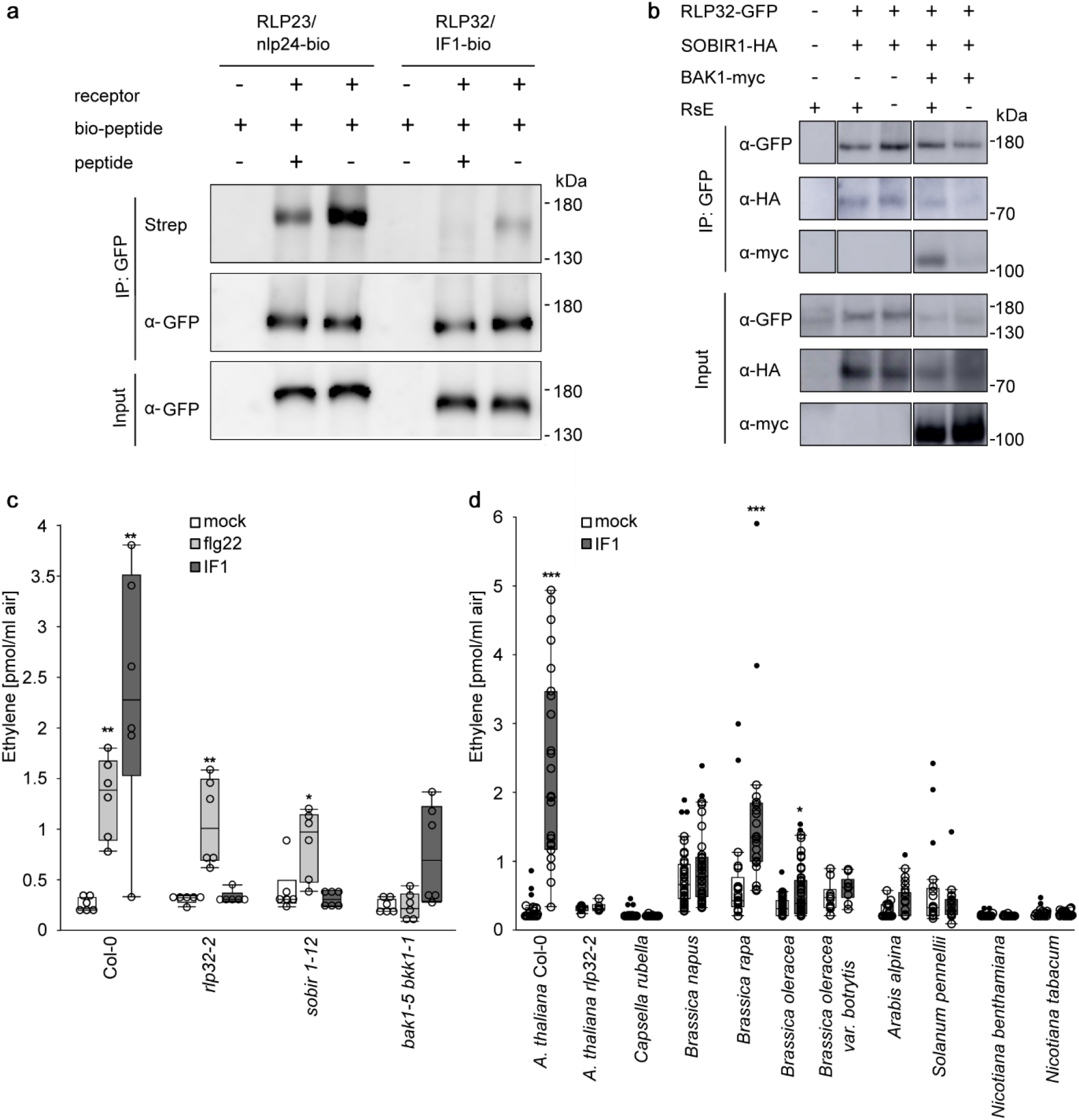
RLP32 binds to IF1 and forms complexes with SOBIR1 and BAK1. **a**, Protein blot analysis of crosslinking assays using GFP-trap purified proteins obtained from RLP32-GFP transgenic *N. benthamiana* plants (input) treated with 30 nM biotinylated IF1 (IF-bio) as ligand. A 1000-fold molar excess of unlabeled peptide was used as competitor of ligand binding. Transiently expressed RLP23-GFP and biotinylated nlp24 (nlp24-bio) served as controls. **b**, RLP32-GFP, BAK1-myc and SOBIR1-HA proteins were transiently expressed in *N. benthamiana* and treated with RsE (+) as indicated. Protein extracts (input) were subjected to co-immunoprecipitation using GFP-trap beads (IP: GFP), and bound proteins were analyzed by protein blotting using tag-specific antisera. **c**, Ethylene accumulation in *Arabidopsis* Col-0 wild-type plants or indicated mutants after treatment with IF1 or flg22. Bars represent mean values ± SD (*n*=6, pooled from two experiments, * *P*≤0,5, ** *P*≤0,01, *** *P*≤0,001, Steel’s test with mock treatment as a control) **d**, Ethylene accumulation in indicated plant species after treatment with IF1. Bars represent mean values ± SD (*n*≥6, pooled from three experiments, * *P*≤0,5, ** *P*≤0,01, *** *P*≤0,001, Mann-Whitney-U-test). Experiments were performed at least twice, with similar results.

LRR-RP-type PRRs constitutively interact with SOBIR1 and recruit the co-receptor BAK1 into a receptor-ligand complex in a ligand-dependent manner. We have conducted co-immunoprecipitation assays in transiently transformed *N. benthamiana* plants to demonstrate IF1-induced formation of RLP32-SOBIR1-BAK1 complexes. As shown in Fig. 6b, RLP32 SOBIR1 complexes were formed independently of IF1 treatment. In contrast, RLP32 SOBIR1 BAK1 complexes were formed only in plants treated with IF1 (Fig. 6b). *Arabidopsis sobir1* and *bak1-5/bkk1-1* mutants proved insensitive to IF1-induced ethylene production, thus confirming a role of SOBIR1 and BAK1 in RLP32-mediated plant defense (Fig. 6c).

### IF1 sensitivity is restricted to selected Brassicaceae

To assess the distribution of IF1 recognition systems among plants, we tested IF1-inducible ethylene production in close relatives of *Arabidopsis* (Fig. 6d). *Capsella rubella, Arabis alpina* or *Thelungiella halophila* all lacked IF1 sensitivity, but *Brassica rapa* and *B. oleracea* responded to IF1 treatment. In contrast, a breeding variant of *B. oleracea*, *B. oleracea* var. *botrytis*, proved insensitive to IF1 treatment. Likewise, members of the *Solanaceae* (*Nicotiana benthamiana*, *Nicotiana tabacum*, *Solanum pennellii*) did not recognize IF1 (Fig. 6d). IF1 sensitivity appears to be rarer, or even absent in these species.

## Discussion

Here we report biochemical purification and mass spectrometry-based identification of bacterial translation initiation factor 1 (IF1), we characterize its immunogenic activity in *Arabidopsis* and related Brassicaceae species, and we identify RLP32 as an IF1 receptor. IF1, a 8.2 kDa polypeptide, is likely one component within a low molecular weight protein fraction from *R. solanacearum* that was previously shown to trigger immunity in *Arabidopsis* in an FLS2-independent manner^29^. IF1 molecules from taxonomically unrelated proteobacteria are active inducers of *Arabidopsis* defense (Fig. 2d, Supplementary Fig. 13), which is in agreement with their high primary and tertiary structure conservation. Generally, IF1 conforms to the definition of classical PAMPs as immunogenic molecules that are (i) ubiquitous to whole classes of microbes, (ii) structurally conserved across microbial species or genus boundaries, and (iii) not found in potential host organisms.

Bacterial IF1 and translation elongation factor EF-Tu constitute not only different parts of the same molecular machineries, but share similar immunogenic activities in *Arabidopsis* (Fig. 4e). During plant infection, both patterns are likely to trigger immune defenses simultaneously. However, concomitant application of both patterns at non-saturating concentrations did not result in additive or synergistic increases in plant defense output (Fig. 4e), suggesting that individual PRR systems may act rather independently and as entirely redundant entities. In other words, plants may integrate various external stimuli to mount an appropriate output response, suggesting that signal input may not necessarily be directly proportionally linked to plant defense outputs.

Immunogenic activities of virtually all known microbe-derived plant defense elicitors can be ascribed to small epitopes within these molecules^1^. IF1 appears to be a remarkable exception to this rule as our collective experimental efforts have revealed that the IF1 tertiary fold may be required for its elicitor activity. Testing a library of nested peptides spanning intact IF1 or of peptides covering individual IF1 secondary structure motifs failed to reveal a small peptide elicitor (Fig. 4a,b). Likewise, peptide mixes and larger terminal deletion mutants affecting IF1 secondary structure motifs lacked elicitor activity (Fig. 4a,b). IF1 shares with elicitor-inactive bacterial cold shock protein CspA a highly conserved 5-strand β-barrel fold^31^, and carries an additional short α-helical motif between β-strands 3 and 4. However, introduction of single or higher order mutations into this helical motif did not affect IF1 elicitor activity, nor did engineering of the IF1 helix into CspA (CspA-IF1 Helix) result in an active elicitor (Fig. 4c). Together with the observed heat instability of IF1 activity (Fig. 4d), our data suggest that tertiary structure features rather than primary sequence motifs determine IF1 elicitor activity. While apparently uncommon for plant immunogenic patterns, structural fold requirements for the activation of pattern-induced immunity have been reported from metazoans. For example, activation of human TOLL-LIKE RECEPTOR 5 (TLR5) is brought about by recognition of large internal helical structures within intact flagellin^33^.

We have exploited natural variation within *Arabidopsis* accessions to identify the IF1 receptor. A continuous spectrum of phenotypic variation of IF1 sensitivity limited the power of GWAS (Genome-Wide Association Studies) studies (Supplementary Fig. 3). The use of heterogeneous RsE fractions, which could potentially contain multiple elicitors, further increased the risk of ambiguous phenotype determination in segregating populations. These hindrances would also prove challenging to classical PCR polymorphism marker-based mapping approaches that are low-throughput and thus require repetitive phenotyping. To avoid laborious marker testing and phenotyping errors inherent to map-based cloning, we employed a genotyping-by-sequencing (GBS) approach to identify and score approximately 16,000 markers in a non-reference bi-parental population. QTL analysis of the F2 segregating population derived of representative extreme morphs enabled the identification of a sizable genomic interval in chromosome 3 encoding RsE sensitivity (Fig. 1c, Supplementary Figs. 5 and 6). Subsequent RAD-seq-based identification of recombination break boundaries from the very same sample set narrowed down the RsE-encoding region to a 1.1 Mb fragment encoding a small number of receptor candidates (Fig. 1d). We propose that RAD-seq-based QTL mapping is superior to classical mapping approaches as linking particular phenotypes to defined genomic loci is supported by strong statistical analyses due to the use of large markers sets^34^.

Reverse genetics analysis identified RLP32 as a locus conferring RsE and IF1 sensitivity. Subsequent genetic and biochemical assays established a role of RLP32 as the IF1 receptor. Evidence for this is based on the following findings: (1) *rlp32* mutants do not mount IF1-inducible defenses, (2) ectopic expression of RLP32 in IF1-insensitive *Arabidopsis* accessions and in *rlp32* mutants confers IF1 sensitivity, (3) production of RLP32 in IF1-insensitive *N. benthamiana* confers IF1 sensitivity, (4) RLP32 specifically binds IF1, (5) RLP32 forms with SOBIR1 and BAK1 a ternary immune receptor complex similar to that known for other *Arabidopsis* LRR-RP-type PRRs, (6) wild-type plants, but not *rlp32* mutants are resistant to bacterial infection following pre-treatment with IF1.

Distribution of LRR-RP-type PRRs is remarkably restricted to individual plant genera. Functional homologs of *Arabidopsis* RLP1, RLP23 and RLP42 are only found in this genus as well as in a few related Brassicaceae^17,35,36^. Likewise, a tomato receptor for fungal xylanase (EIX2) or *N. benthamiana* sensors for CspA (CSPR) and a fungal hydrolase (RXEG1) exhibit genus-specific distribution^21,22,37,38^. RLP32 is no exception to this rule as IF1 sensitivity appears rare outside Brassicaceae (Fig. 6d) and shows significant within-genus variation (Supplementary Fig. 3). In conclusion, a rather genus-specific distribution as well as significant accession-specific sequence polymorphisms among *Arabidopsis* PRRs suggest highly dynamic evolution of PTI sensors in this plant.

Identification of the ligand-receptor pair IF1/RLP32 illustrates that recognition of a single pathogen species by a given host plant can be enormously complex. In addition to RLP32, *Arabidopsis* has evolved at least five additional receptor systems to sense the bacterial pathogen, *P. syringae*. These receptors comprise well-studied FLS2 and EFR as well as sensors for bacterial mc-3-OH-fatty acids (LORE), peptidoglycan (LYM1-LYM3-CERK1) and xanthine/uracil permease (XPS1)^16,18,19,23,24^. We can only speculate why a single plant species may have evolved such a number of redundant microbial sensing systems. It seems to be reasonable to assume that plant PRR complexity may not be brought about by co-evolution of these two organisms alone, but may result from multiple independent evolutionary processes that were driven by exposure to various microbial threats. Redundancy in *Arabidopsis* PRRs may explain, however, accession-specific losses of recognition specificities that have been reported for most microbial sensor systems in this plant.

## Materials and Methods

### Plant materials and growth conditions

All plants except for *Solanaceae* were grown on soil for 5-6 weeks under standard conditions (150 μmol/cm^2^s light for 8 h, 40-60 % humidity, 22°C). *Arabidopsis* accession Col-0 was the background for all mutants used in this study: *bak1-5 bkk1-1*^39^, *fls2 efr*^40^, *rlp32-2* (SM_3_33092, N119803), *rlp32-3* (SALK_137467C, N657024), *rlp32-4* (SM_3_33695, N120406), *rlp32-5* (SM3_15851, N106446), *rlp31-1*^41^, *rlp31-2*^41^, *rlp33-2*^41^, *rlp33-3*^41^, *sobir1-12*^42^, SALK_143696 (N643696, At3g05990), SALK_203784C (N692234, At3g05990), SALK_146601C (N660645, At3g07040), SAIL_918_H07 (N879828, At3g07040). Seeds of T-DNA or transposon insertion lines were purchased from ABRC or NASC, respectively. Insertions of these alleles were confirmed by comparing flanking sequences according to the Col-0 reference genome. For *rlp32* mutants, insertions were additionally verified by flanking fragment sequencing (Supplementary Fig. 7) using the Phire Plant Direct PCR Kit (Thermo Fisher Scientific) and primers listed in Supplementary Table 1. *Arabidopsis* natural germplasms used in this study are part of the 80 re-sequenced accessions (ABRC CS76427)^43^ and the Nordborg collection^44^. *N. benthamiana, N. tabacum* and *Solanum pennellii* plants were grown for 4-5 weeks on soil in the greenhouse (16 h light, 60-70 % humidity, 22 °C). For stable transformation, *N. benthamiana* plants were grown for 7-8 weeks in sterile culture on MS medium containing 2 % sucrose (13 h light, 23 °C). *Arabidopsis* and *N. benthamiana* plants used for bacterial infection assays were grown under a translucent cover on soil under standard conditions (150 μmol/cm^2^s light for 8 h, 40-60 % humidity, 22°C).

### Elicitors used in this study

Synthetic IF1 protein and all peptides, as well as nlp20^45^, nlp24-bio^17^, flg22^46^ and csp22^32^ were purchased from Genscript Inc. (Piscataway, New Jersey, US) and were dissolved in 100 % DMSO as 10 mM stock solution, flg22 was dissolved in 0,1 % BSA, 0,1 M NaCl as 1 μM stock solution, full-length IF1 was dissolved in sterile filtered 10 mM MES pH 5.7 or in 100 % DMSO as a 100 μM stock solution, IF1 A1-S36 was dissolved in 3 % ammonium water as a 1 mM stock solution, IF1 G37-R71 and IF1 N27-R71 were dissolved in water as 1 mM stock solutions. All working dilutions were prepared in water prior to use. If not stated otherwise, synthetic elicitors used in this study were applied at 1 μM concentrations. *Penicillium*-derived elicitor PEN and SCFE1 were used at concentrations of 90 μg/ml and 0.25 μg/ml, respectively^30^. As elicitor activities and protein contents varied among RsE-containing fractions, sample volumina ranging from 1-20 μl were used to detect RsE elicitor activities.

### RsE preparation and LC-MS/MS analysis

RsE purification was essentially performed as previously described^34^. In brief, *Ralstonia solanacearum* GMI1000^47^ was cultivated in Kelman medium (10 g/l glucose, 10g/l peptone, 1 g/l casein hydrolysate, pH 6.5)^48^ at 28°C for 36 to 48 h at 200 rpm. 5 l cell culture was boiled, cooled on ice and centrifuged at 5000 g for 15 min. For protein precipitation, ammonium sulfate was added to the supernatant to a 90 % saturation and precipitated proteins were collected by centrifugation at 10,000 g and then re-dissolved in 50 mM MES, pH 5.2. The crude extract was dialyzed overnight at 4°C in 50 mM MES, pH 5.2 (ZelluTrans, Carl Roth, or Slide-A-Lyzer Dialysis Cassette, Thermo Fisher Scientific, each 3.5 kDa) before loading onto a cation exchange HiTrapSP FF column (GE Healthcare, Uppsala, Sweden) using an Äkta explorer system (GE Healthcare, loading speed 1 ml/min) with buffers A (50 mM MES, pH 5.2) and B (50 mM MES, 0.5 M KCl, pH 5.2). Bound proteins were eluted with 100 % buffer B, and pooled fractions were dialyzed in 50 mM Tris/HCl, pH 8.5. Anion exchange chromatography on dialyzed extracts was performed using an HiTrapQ FF column (GE Healthcare) with buffer C (50 mM Tris/HCl, pH 8.5) and D (50 mM Tris/HCl, 0.5 M KCl, pH 8.5). The flow-through was dialyzed in 50 mM MES, pH 5.2 before loading onto a Source 15S 4.6/100 PE cation exchange column (GE Healthcare) for high resolution protein separation at a loading speed of 1 ml/min, followed by gradient elution with buffer B. Eluted fractions were tested for ethylene-inducing activity in *Arabidopsis fls2 efr* mutant plants and active fractions were pooled and stored at −20 °C.

For LC-MS/MS analysis active fractions purified via cation exchange chromatography were further purified by reverse phase high pressure liquid chromatography (HPLC) using a C8 column ZORBAX 300SB, 5μm, 4.6 × 150 mm, (Agilent, Waldbronn, Germany) and a gradient of buffers E (H_2_O, 0.1 % TFA) and F (Isopropanol, 0.1 % TFA) at a flow rate of 1 ml/min. Fractions containing ethylene-inducing activity were analyzed by LC-MS/MS as described^17^.

### Rad-Seq and QTL analysis

Restriction site–associated DNA sequencing (RAD-seq) and quantitative trait locus mapping using package R/qtl were conducted using the F_2_ mapping population of a *Arabidopsis* ICE153 × ICE73 cross as previously described^34^.

### IF1 synthesis and recombinant expression

*In vitro* transcription and translation (TnT) was performed with full length IF1 cloned into pET-pDEST42 (Thermo Fisher Scientific, for primer sequences see Supplementary Table 2) and the TnT® Quick Coupled Transcription/Translation System (for T7 promoter, Promega) according to the manufacturers protocol. As a control the provided Luciferase construct was used.

For recombinant protein expression in *E. coli* BL21AI, full length IF1 from different bacteria or IF1-fragments I6-R71 and A1-I66 were cloned into pET-pDEST42 (Thermo Fisher Scientific, for primer sequences see Supplementary Table 2). Protein expression was induced by 0.2 % L-arabinose and 1 mM IPTG for 24 hours at 17°C and 220 rpm. The cell pellets were resuspended in binding buffer A containing 20 mM KP_i_, pH 7.4, 500 mM KCl and 50 mM imidazole. After sonication and centrifugation (45 min, 14,000 rpm, 4 °C), the clear supernatant was applied to a 1 ml HisTrapFF column (GE Healthcare). C-terminally 6xHis-tagged proteins were collected in 1 ml fractions with an elution buffer gradient (1 ml/min, 0-100% buffer B: 20 mM KP_i_, pH 7.4, 500 mM KCl, 500 mM imidazole). The protein concentration was determined according to Bradford^49^ using the Roti-Quant solution (Carl Roth).

For recombinant protein expression in *Pichia pastoris* GS115 (Multi-Copy Pichia Expression Kit Instructions, Thermo Fisher Scientific), constructs of IF1, mutant versions of IF1, CspA with IF1 helix or CspA in the secretory expression plasmid pPICZalphaA were generated. IF1 wild type coding sequence was amplified from the *E. coli* expression construct. IF1 mutant constructs were generated using the GeneArt Site-Directed Mutagenesis System (Thermo Fisher Scientific) and AccuPrime Pfx DNA Polymerase (Thermo Fisher Scientific). CspA and CspA-IF1-helix coding sequences were amplified from synthetic gene constructs (Eurofins). Protein purification from *P. pastoris* culture medium was achieved by affinity chromatography on HisTrap excel columns (equilibrated in 20 mM KP_i_ pH 7.4, 500 mM KCl). Following washing (20 mM KP_i_ pH 7.4, 500 mM KCl, 20 mM imidazole) and elution (buffer gradient 0–500 mM imidazole in equilibration buffer), IF1 or CspA containing fractions were pooled and dialyzed against H_2_O. Protein concentrations were calculated by UV spectroscopy (wavelength λ280) using the protparam tool (http://web.expasy.org/protparam) to determine protein-specific extinction coefficients ε280 for each protein. Determinations were verified by SDS-PAGE using a standard protein solution.

### Generation of transgenic plants

*RLP32* coding sequence with or without the native promoter were amplified with Pfu DNA Polymerase (Thermo Fisher Scientific) using primers listed in Supplementary Table 2, and cloned into the pCR®8/GW/TOPO®-TA vector (Thermo Fisher Scientific). For 35S-promoter driven expression in *N. benthamiana*, *RLP32* coding sequence was fused to a C-terminal GFP-tag in pB7FWG2.0^50^, for native-promoter-driven expression in *Arabidopsis*, *RLP32* promoter and coding sequence were recombined into pGWB1 (no tag) or pGBW4 (C-terminal GFP-tag)^51^, respectively. Transient and stable transformation of *Arabidopsis* and *N. benthamiana* was performed as described previously^17^.

### Immune assays and bacterial infections

The determination of ethylene accumulation, the production of reactive oxygen species (ROS), the staining of callose appositions, the detection of activated MAPKs and the histochemical detection of GUS enzyme activity in *pPR-1::GUS* transgenic plants were performed as previously described^17,30^. In *Arabidopsis* plants, a 24 h-priming using 1 μM nlp20 or IF1 and subsequent infection with a final cell density of 10^4^ cfu/ml *Pseudomonas syringae* pv. *tomato* DC3000 (*Pst* DC3000) was performed as described^17^. *N. benthamiana* plants were likewise infected with *P. syringae* pv. *tomato* DC3000 hrcC-^52^ at a final cell density of 2*10^4^ cfu/ml and harvested after 0 and 4 days.

### Immunoprecipitation assays and *in vivo* cross-linking

For co-immunoprecipitation, RLP32-GFP (in pB7FWG2^50^) and co-receptors SOBIR1-HA (pGWB14^51^) and BAK1-myc (pGWB17^51^) were transiently expressed in *N. benthamiana*, and leaf material was harvested 5 min after infiltration of RsE or 10 mM MgCl_2_ as negative control. 200 mg ground leaf material was subjected to protein extraction and immunoprecipitation using GFP-Trap beads (ChromoTek, IZB Martinsried, Germany) as previously described^17^.

*In vivo* cross-linking experiments were essentially done as described^17^ using leaves of *N. benthamiana* stably expressing *p35S::RLP32–GFP* (in pB7FWG2^50^), or transiently expressing *p35S::RLP23–GFP*^17^. In brief, leaves were infiltrated with 30 nM biotinylated IF1, 30 nM biotinylated nlp24 or 10 mM MgCl_2_, respectively, with or without 30 μM unlabeled synthesized IF1 as competitor. Five min after peptide treatment, 2 mM ethylene glycol bis(succinimidyl succinate) (EGS) was infiltrated into the same leaves and leaf samples were harvested after further 15 min. 300 mg of the sample was used for protein extraction and immuno-adsorption to GFP-Trap beads as described^17,53^.

Protein blots were probed either directly with a Streptavidin-alkaline phosphatase conjugate (Roche) or with antibodies raised against GFP (Sicgen, Torrey Pines Biolabs), HA- or Myc-tags (Sigma-Aldrich) followed by staining with secondary antibodies coupled to alkaline phosphatase (Sigma-Aldrich) and CDP-Star (Roche) as substrate. Chemiluminescence was detected using a CCD-camera (Viber Louromat, PeqLAB).

### Statistical analysis

All statistical analyses were carried out with SAS jmp. All normal distribution data sets of the infection assays were evaluated using the post-hoc comparisons following one-way ANOVA (Dunnett’s test with control) multiple comparison analysis at a probability level of *P* < 0.05. EC_50_ values and curve fit were calculated using 4P Rodbard Model comparison (three parametric logistic regression). All data sets with no normal distribution were evaluated with nonparametric tests: for comparison of two data sets Mann-Whitney-U-test was used; for multiple comparison analysis either Steel-test with control or Steel-Dwass-test were used, as indicated in the figure legends.

## Acknowledgements

Research in the lab of T.N. was funded by DFG grants Nu70/9-1, Nu70/9-2, Nu 70/15-1, Nu70/16-1 and Nu70/17-1 and SFB1101.

## Author contributions

L.F., K.F., E.M., I.A., E.C. and T.N. conceived and designed the experiments; L.F., K.F., E.M., I.A., R.N.P. and L.Z. conducted experiments; L.F., K.F., E.M., I.A., R.N.P., L.Z., M.A., S-T.K., E.C., D.W., A.A.G. and T.N. analysed data; and R.N.P., E.C., A.A.G. and T.N. wrote the manuscript. All authors discussed the results and commented on the manuscript.

## Additional information

Supplementary files include Supplementary Fig.1-15 and Supplementary Tables 1-2.

## References

1 Albert, I., Hua, C., Nürnberger, T., Pruitt, R. N. & Zhang, L. Surface sensor systems in plant immunity. Plant Physiol. 182, 1582–1596 (2020).

2 Wan, W. L., Fröhlich, K., Pruitt, R. N., Nürnberger, T. & Zhang, L. Plant cell surface immune receptor complex signaling. Curr. Opin. Plant Biol. 50, 18–28 (2019).

3 Macho, A. P. & Zipfel, C. Plant PRRs and the activation of innate immune signaling. Mol. Cell 54, 263–272 (2014).

4 Zhou, J. M. & Zhang, Y. Plant immunity: danger perception and signaling. Cell 181, 978–989 (2020).

5 Dodds, P. N. & Rathjen, J. P. Plant immunity: towards an integrated view of plant-pathogen interactions. Nat. Rev. Genet. 11, 539–548 (2010).

6 Thomma, B. P., Nürnberger, T. & Joosten, M. H. Of PAMPs and effectors: the blurred PTI-ETI dichotomy. Plant Cell 23, 4–15 (2011).

7 Ngou, B. P. M., Ahn, H.-K., Ding, P. & Jones, J. D. Mutual potentiation of plant immunity by cell-surface and intracellular receptors. bioRxiv, 2020.2004.2010.034173 (2020).

8 Yuan, M. et al. Pattern-recognition receptors are required for NLR-mediated plant immunity. bioRxiv, 2020.2004.2010.031294 (2020).

9 Pruitt, R. N. et al. *Arabidopsis* cell surface LRR immune receptor signaling through the EDS1-PAD4-ADR1 node. bioRxiv, 2020.2011.2023.391516 (2020).

10 Böhm, H., Albert, I., Fan, L., Reinhard, A. & Nürnberger, T. Immune receptor complexes at the plant cell surface. Curr. Opin. Plant Biol. 20, 47–54 (2014).

11 Chinchilla, D., Shan, L., He, P., de Vries, S. & Kemmerling, B. One for all: the receptor-associated kinase BAK1. Trends Plant Sci. 14, 535–541 (2009).

12 Wan, W. L. et al. Comparing Arabidopsis receptor kinase and receptor protein-mediated immune signaling reveals BIK1-dependent differences. New Phytol. 221, 2080–2095 (2019).

13 Hegenauer, V. et al. Detection of the plant parasite Cuscuta reflexa by a tomato cell surface receptor. Science 353, 478–481 (2016).

14 Hegenauer, V. et al. The tomato receptor CuRe1 senses a cell wall protein to identify Cuscuta as a pathogen. Nat. Commun. 11, 5299 (2020).

15 Steinbrenner, A. D. et al. A receptor-like protein mediates plant immune responses to herbivore-associated molecular patterns. Proc. Natl. Acad. Sci. USA (2020).

16 Gomez-Gomez, L. & Boller, T. FLS2: an LRR receptor-like kinase involved in the perception of the bacterial elicitor flagellin in Arabidopsis. Mol. Cell 5, 1003–1011 (2000).

17 Albert, I. et al. An RLP23-SOBIR1-BAK1 complex mediates NLP-triggered immunity. Nat. Plants 1, 15140 (2015).

18 Zipfel, C. et al. Perception of the bacterial PAMP EF-Tu by the receptor EFR restricts *Agrobacterium*-mediated transformation. Cell 125, 749–760 (2006).

19 Mott, G. A. et al. Genomic screens identify a new phytobacterial microbe-associated molecular pattern and the cognate Arabidopsis receptor-like kinase that mediates its immune elicitation. Genome Biol. 17, 98 (2016).

20 Hind, S. R. et al. Tomato receptor FLAGELLIN-SENSING 3 binds flgII-28 and activates the plant immune system. Nat. Plants 2, 16128 (2016).

21 Wang, L. et al. The pattern-recognition receptor CORE of *Solanaceae* detects bacterial cold-shock protein. Nat. Plants 2, 16185 (2016).

22 Saur, I. M. et al. NbCSPR underlies age-dependent immune responses to bacterial cold shock protein in *Nicotiana benthamiana*. Proc. Natl. Acad. Sci. USA 113, 3389–3394 (2016).

23 Kutschera, A. et al. Bacterial medium-chain 3-hydroxy fatty acid metabolites trigger immunity in *Arabidopsis* plants. Science 364, 178–181 (2019).

24 Willmann, R. et al. *Arabidopsis* lysin-motif proteins LYM1 LYM3 CERK1 mediate bacterial peptidoglycan sensing and immunity to bacterial infection. Proc. Natl. Acad. Sci. USA 108, 19824–19829 (2011).

25 Fischer, I., Dievart, A., Droc, G., Dufayard, J. F. & Chantret, N. Evolutionary dynamics of the leucine-rich repeat receptor-like kinase (LRR-RLK) subfamily in Angiosperms. Plant Physiol. 170, 1595–1610 (2016).

26 Shiu, S. H. & Bleecker, A. B. Expansion of the receptor-like kinase/Pelle gene family and receptor-like proteins in *Arabidopsis*. Plant Physiol. 132, 530–543 (2003).

27 Shiu, S. H. et al. Comparative analysis of the receptor-like kinase family in *Arabidopsis* and rice. Plant Cell 16, 1220–1234 (2004).

28 Boutrot, F. & Zipfel, C. Function, discovery, and exploitation of plant pattern recognition Receptors for Broad-Spectrum Disease Resistance. Annu. Rev. Phytopathol. 55, 257–286 (2017).

29 Pfund, C. et al. Flagellin is not a major defense elicitor in *Ralstonia solanacearum* cells or extracts applied to *Arabidopsis thaliana*. Mol. Plant Microbe Interact. 17, 696–706 (2004).

30 Zhang, W. et al. *Arabidopsis* receptor-like protein30 and receptor-like kinase suppressor of BIR1-1/EVERSHED mediate innate immunity to necrotrophic fungi. Plant Cell 25, 4227–4241 (2013).

31 Sette, M. et al. The structure of the translational initiation factor IF1 from *E-coli* contains an oligomer-binding motif. Embo J. 16, 1436–1443 (1997).

32 Felix, G. & Boller, T. Molecular sensing of bacteria in plants. The highly conserved RNA-binding motif RNP-1 of bacterial cold shock proteins is recognized as an elicitor signal in tobacco. J. Biol. Chem. 278, 6201–6208 (2003).

33 Smith, K. D. et al. Toll-like receptor 5 recognizes a conserved site on flagellin required for protofilament formation and bacterial motility. Nat. Immunol. 4, 1247–1253 (2003).

34 Fan, L., Chae, E., Gust, A. A. & Nürnberger, T. Isolation of novel MAMP-like activities and identification of cognate pattern recognition receptors in *Arabidopsis thaliana* using next-generation sequencing (NGS)-based mapping. Curr. Protoc. Plant Biol. 2, 173–189 (2017).

35 Jehle, A. K. et al. The receptor-like protein ReMAX of *Arabidopsis* detects the microbe-associated molecular pattern eMax from *Xanthomonas*. Plant Cell 25, 2330–2340 (2013).

36 Zhang, L. et al. Fungal endopolygalacturonases are recognized as microbe-associated molecular patterns by the *Arabidopsis* receptor-like protein RESPONSIVENESS TO BOTRYTIS POLYGALACTURONASES1. Plant Physiol. 164, 352–364 (2014).

37 Wang, Y. et al. Leucine-rich repeat receptor-like gene screen reveals that *Nicotiana* RXEG1 regulates glycoside hydrolase 12 MAMP detection. Nat. Commun. 9, 594 (2018).

38 Ron, M. & Avni, A. The receptor for the fungal elicitor ethylene-inducing xylanase is a member of a resistance-like gene family in tomato. Plant Cell 16, 1604–1615 (2004).

39 Schwessinger, B. et al. Phosphorylation-dependent differential regulation of plant growth, cell death, and innate immunity by the regulatory receptor-like kinase BAK1. PLoS Genet. 7, e1002046 (2011).

40 Nekrasov, V. et al. Control of the pattern-recognition receptor EFR by an ER protein complex in plant immunity. Embo J. 28, 3428–3438 (2009).

41 Wang, G. et al. A genome-wide functional investigation into the roles of receptor-like proteins in *Arabidopsis*. Plant Physiol. 147, 503–517 (2008).

42 Gao, M. et al. Regulation of cell death and innate immunity by two receptor-like kinases in *Arabidopsis*. Cell Host Microbe 6, 34–44 (2009).

43 Cao, J. et al. Whole-genome sequencing of multiple *Arabidopsis thaliana* populations. Nat. Genet. 43, 956–963 (2011).

44 Nordborg, M. et al. The pattern of polymorphism in *Arabidopsis thaliana*. PLoS Biol. 3, 1289–1299 (2005).

45 Böhm, H. et al. A conserved peptide pattern from a widespread microbial virulence factor triggers pattern-induced immunity in *Arabidopsis*. PLoS Pathog. 10, e1004491 (2014).

46 Felix, G., Duran, J. D., Volko, S. & Boller, T. Plants have a sensitive perception system for the most conserved domain of bacterial flagellin. Plant J. 18, 265–276 (1999).

47 Salanoubat, M. et al. Genome sequence of the plant pathogen *Ralstonia solanacearum*. Nature 415, 497–502 (2002).

48 Kelman, A. The relationship of pathogenicity in *Pseudomonas solanacearum* to colony appearance on a tetrazolium medium. Phytopathology 44, 693–695 (1954).

49 Bradford, M. M. A rapid and sensitive method for the quantitation of microgram quantities of protein utilizing the principle of protein-dye binding. Anal. Biochem. 72, 248–254 (1976).

50 Karimi, M., De Meyer, B. & Hilson, P. Modular cloning in plant cells. Trends Plant Sci. 10, 103–105 (2005).

51 Nakagawa, T. et al. Development of series of gateway binary vectors, pGWBs, for realizing efficient construction of fusion genes for plant transformation. J. Biosci. Bioengineering 104, 34–41 (2007).

52 Boch, J. et al. Identification of *Pseudomonas syringae* pv*. tomato* genes induced during infection of *Arabidopsis thaliana*. Mol. Microbiol. 44, 73–88 (2002).

53 Chinchilla, D. et al. A flagellin-induced complex of the receptor FLS2 and BAK1 initiates plant defence. Nature 448, 497–500 (2007).

