## Supplementary Information for "Genotyping-by-sequencing-based identification of *Arabidopsis* pattern recognition receptor RLP32 recognizing proteobacterial translation initiation factor IF1"

**Supplemental figures**

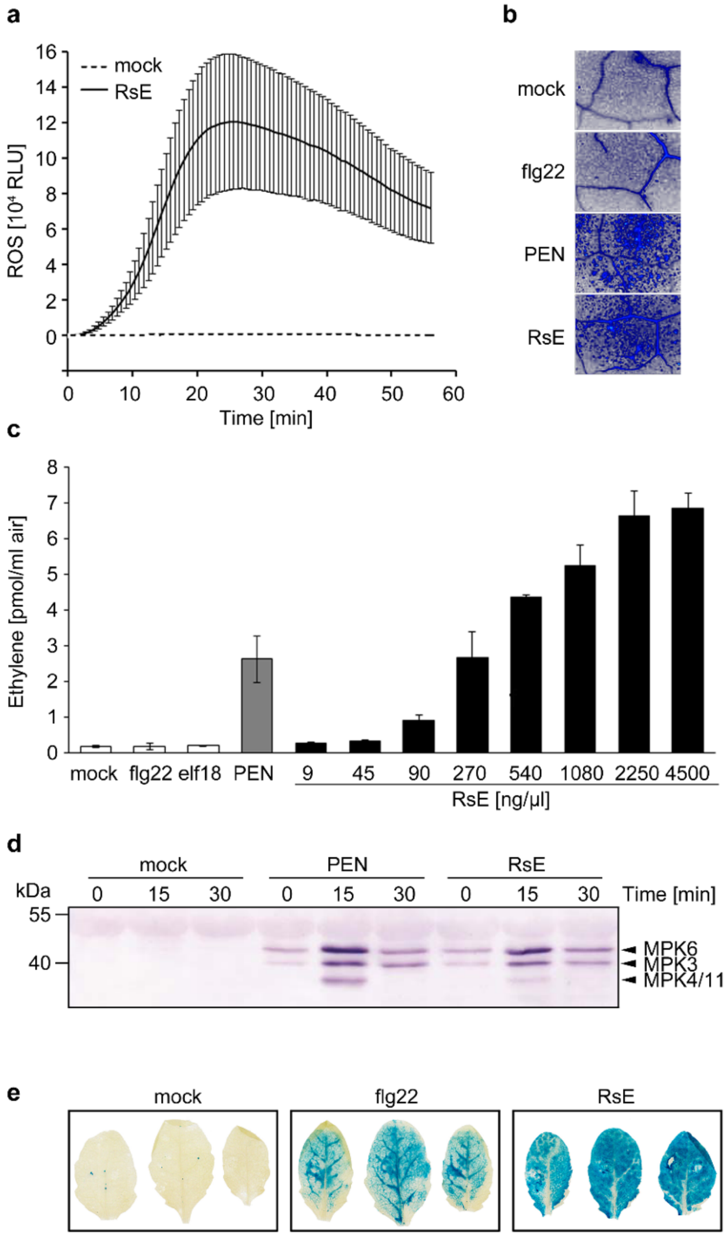


**Supplementary Figure 1. RsE induces plant immune responses in *Arabidopsis*. a,** ROS accumulation in leaf pieces of *Arabidopsis fls2 efr* plants treated with water (mock), or RsE. Given are relative light units (RLU) ± SD (*n* = 6). **b**, Aniline blue stain of callose appositions 24 h after treatment of *fls2 efr* leaves with water (mock), flg22, PEN or RsE, **c,** Ethylene production in *Arabidopsis fls2 efr* plants treated with increasing RsE concentrations. Water treatment (mock) or treatment with flg22, elf18 or PEN served as controls. Bars represent means ± SD of two replicates. **d,** *Arabidopsis fls2 efr* plants were treated for the times indicated with water (mock), PEN or RsE. MAPK activation was detected by immunoblot using phospho-p44/p42 antibodies. **e,** *pPR1::GUS* induction in leaves of three *pPR1::GUS* transgenic *Arabidopsis* lines infiltrated for 24 h with water (mock), flg22 or RsE.

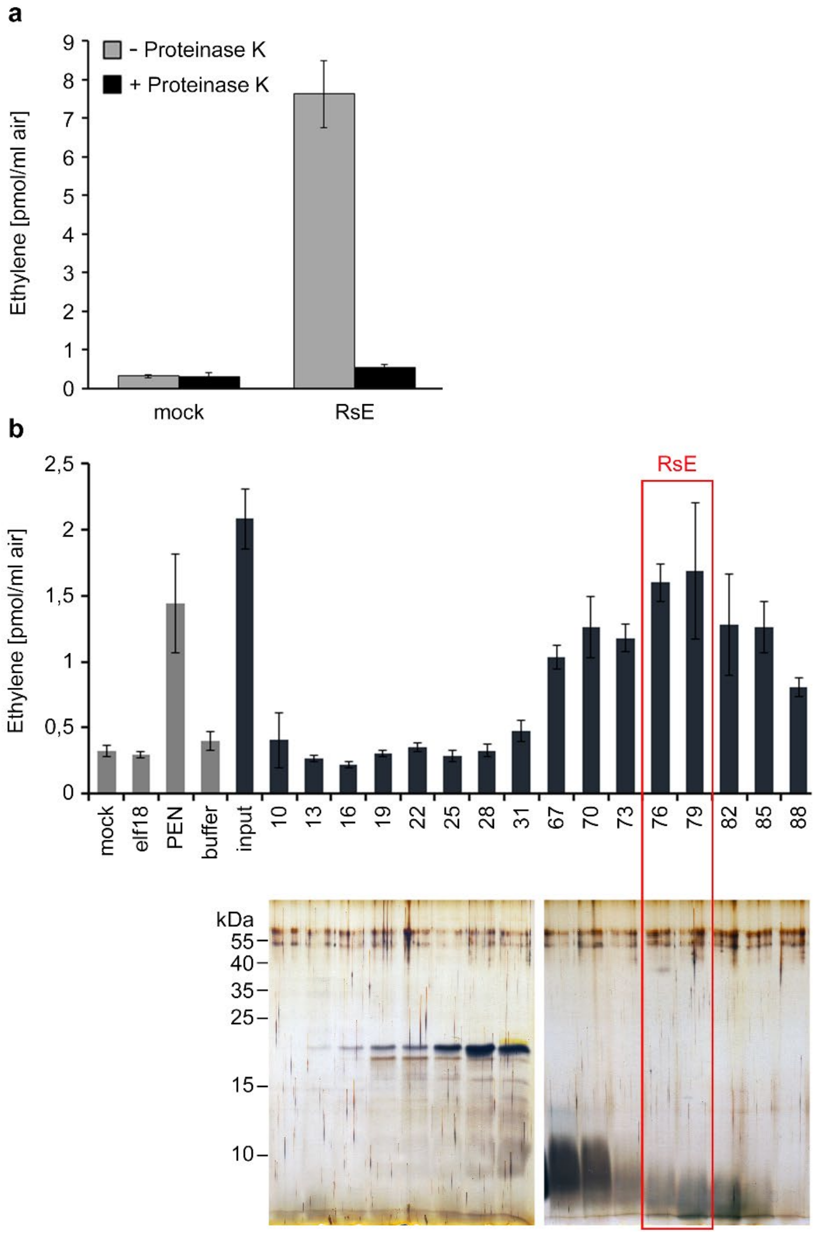


**Supplementary Figure 2. RsE elicitor activity is protease-sensitive and co-migrates with fractions containing low molecular weight proteins. a,** Ethylene accumulation in *Arabidopsis fls2 efr* leaf pieces treated with water (mock) or RsE incubated for 2 h with 0,5 µg/µl Proteinase K (+) or left untreated (-). **b,** Ethylene accumulation (upper panel) in *Arabidopsis fls2 efr* leaf pieces treated with water (mock), elf18, PEN, *R. solanacearum* cell extract (input) or with fractions obtained by gel filtration. Comparison with the elution profile of proteins with known size indicated that highest elicitor activity co-migrates with molecular masses <10 kDa (red box). Gel filtration fractions were analyzed by Tricine-SDS-PAGE followed by silver staining (lower panels).

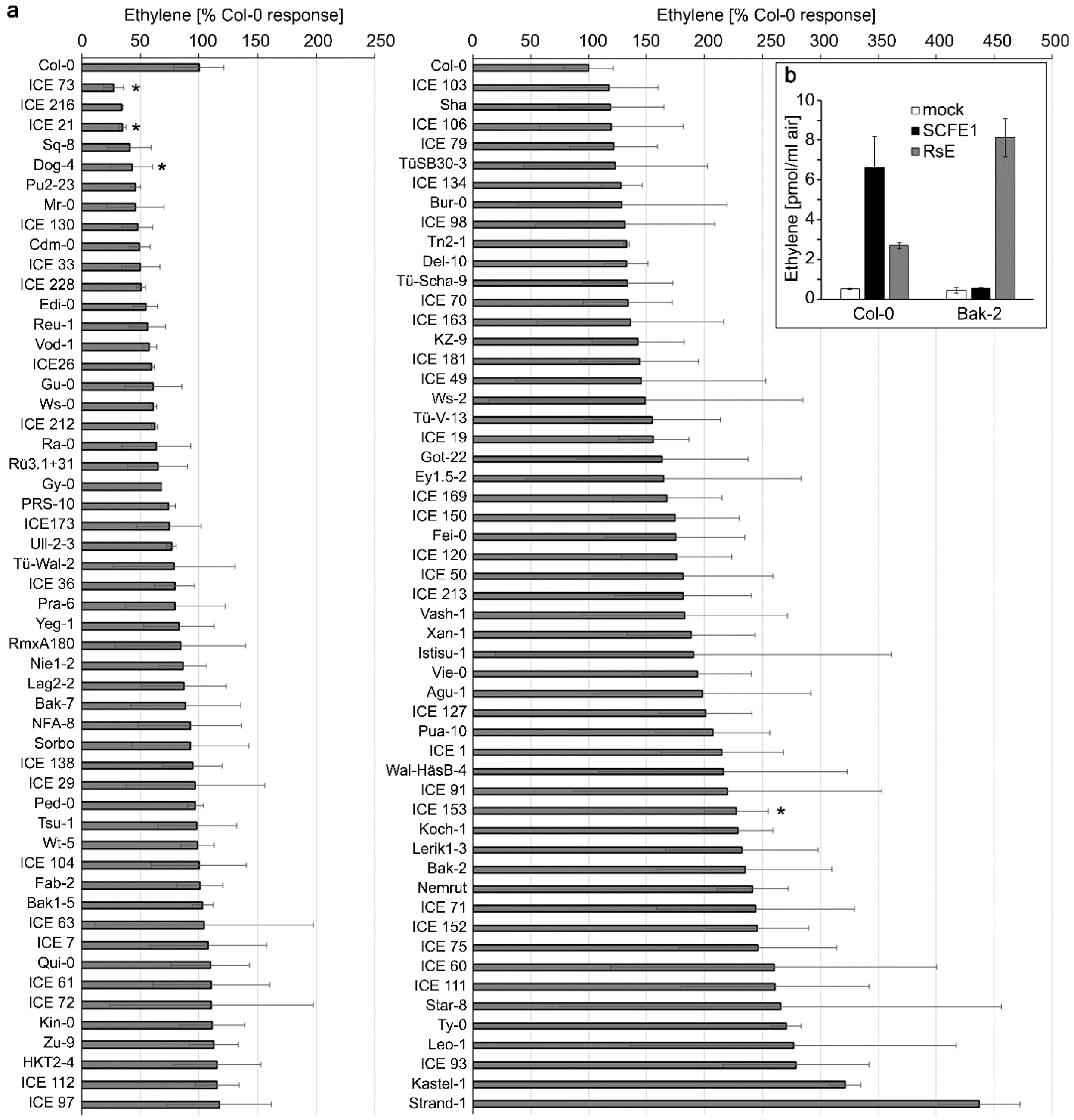


**Supplementary Figure 3. Natural variation in RsE sensitivity among *Arabidopsis* accessions. a**, 106 *Arabidopsis* accessions were tested for ethylene accumulation upon RsE treatment. Results are shown as percentage of the response determined in Col-0. Bars represent means of 2 replicates ± SD, asterisks indicate the accessions used for further analysis. **b**, Ethylene production in *Arabidopsis* Col-0 and Bak-2 plants treated with SCFE1 or RsE. Water treatment (mock) served as control. Bars represent means ± SD of two replicates.

| ♂ \ ♀ | Dog-4 | ICE21 | ICE73 | ICE153 |
|---|---|---|---|---|
| Dog-4 | - | y | y | y |
| ICE21 | y | - | y | y |
| ICE73 | n | y | - | y |
| ICE153 | n | y | y | - |

**Supplementary Figure 4. Summary of crosses generated for allelism tests.** Shown are *Arabidopsis* accessions used for reciprocal crosses between RsE-insensitive (Dog-4, ICE21, ICE73) and RsE-sensitive (ICE153) accessions. "y" indicates that a normal amount of seeds was obtained; "n" indicates failed crosses, "-" indicates crosses not performed.

a

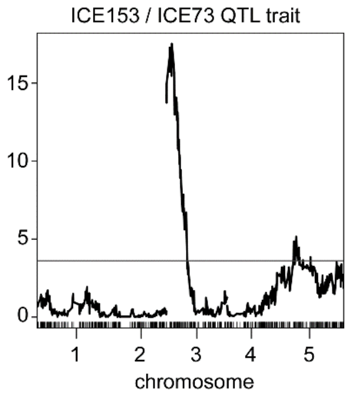


b

| Chr | position | LOD | Chr | position | LOD |
|---|---|---|---|---|---|
| 2 | 18857542 | 0,3023032 | 3 | 3984300 | 19,2518605 |
| 2 | 18858222 | 0,65408905 | 3 | 4005543 | 20,3408255 |
| 2 | 18860115 | 0,63912253 | 3 | 4114238 | 18,4808285 |
| 2 | 19500124 | 0,36913299 | 3 | 4238450 | 18,4739649 |
| 2 | 19511097 | 0,16177605 | 3 | 4303467 | 20,1206988 |
| 2 | 19560723 | 0,35482815 | 3 | 4326887 | 19,2398194 |
| 3 | 20373 | 23,7161535 | 3 | 4415268 | 19,7751107 |
| 3 | 96618 | 20,3589304 | 3 | 4426951 | 18,3413229 |
| 3 | 97041 | 22,1412548 | 3 | 4458826 | 19,3354048 |
| 3 | 99998 | 22,2494445 | 3 | 4786003 | 17,1964815 |
| 3 | 100022 | 21,9456481 | 3 | 4786104 | 18,1941321 |
| 3 | 100053 | 23,1077577 | 3 | 4942846 | 18,8041414 |
| 3 | 1282003 | 24,0596519 | 3 | 5066042 | 15,7032338 |
| 3 | 1286337 | 27,1186078 | 3 | 5091471 | 15,594859 |
| 3 | 1286394 | 24,1720452 | 3 | 5168516 | 18,0124631 |
| 3 | 1321901 | 23,0619188 | 3 | 5225970 | 16,686886 |
| 3 | 1432648 | 26,1204765 | 3 | 5268500 | 12,7434904 |
| 3 | 1546715 | 26,6332744 | 3 | 5533254 | 15,2878901 |
| 3 | 1784868 | 25,4141172 | 3 | 5666710 | 13,7562499 |
| 3 | 1784883 | 25,4103308 | 3 | 5666736 | 13,7562499 |
| 3 | 1784886 | 25,4107724 | 3 | 5940140 | 11,5326217 |
| 3 | 2012983 | 25,010172 | 3 | 6169544 | 10,4838362 |
| 3 | 2063924 | 25,7413858 | 3 | 6169557 | 10,3794269 |
| 3 | 2097697 | 25,9038471 | 3 | 6208231 | 11,0247178 |
| 3 | 2795414 | 26,9539864 | 3 | 6582498 | 11,699298 |
| 3 | 3004962 | 23,5326429 | 3 | 6612203 | 9,29420715 |
| 3 | 3012844 | 23,7849338 | 3 | 6612208 | 9,29407201 |
| 3 | 3012876 | 23,7849338 | 3 | 6612223 | 9,29437913 |
| 3 | 3178197 | 20,2673228 | 3 | 6612255 | 9,49974994 |
| 3 | 3248222 | 20,8794091 | 3 | 6660685 | 9,38376536 |
| 3 | 3326972 | 20,0666037 | 3 | 6702174 | 9,64217411 |
| 3 | 3348485 | 20,8196009 | 3 | 6801332 | 9,0135602 |
| 3 | 3419062 | 20,4317097 | 3 | 7009733 | 7,48710839 |
| 3 | 3833444 | 23,0828747 | 3 | 7437740 | 9,07698364 |
| 3 | 3960217 | 19,6388366 | 3 | 7617984 | 8,83989564 |

**Supplementary Figure 5. rQTL mapping of an RsE-sensitive locus. a,** rQTL mapping for RsE-induced ethylene response in $F_2$ mapping populations of an ICE153 x ICE73 cross. LOD scores from a full genome scan across five chromosomes of Arabidopsis using a QTL trait model for RsE-elicited ethylene scores. The grey lines indicate a genome-wise α equaling to 0.05 LOD thresholds, which defines significant QTLs based on 1,000 permutations. **b,** genomic positions and LOD values derived from rQTL. Given are chromosome number (Chr), position on the chromosome and LOD values. Genomic positions with LOD values above 10 based on rQTL binary trait mapping (see also Figure 1d) are highlighted in grey.

**a**

| Plate 1 | 01 | 02 | 03 | 04 | 05 | 06 | 07 | 08 | 09 | 10 | 11 | 12 |
|---|---|---|---|---|---|---|---|---|---|---|---|---|
| **P1A** | 0 | 0 | 1 | 1 | 1 | 1 | 1 | 1 | 1 | 1 | 1 | 0 |
| **P1B** | 1 | 1 | 0 | 1 | 1 | 1 | 1 | 1 | 1 | 1 | 0 | 0 |
| **P1C** | 1 | 1 | 1 | 1 | 1 | 0 | 0 | 1 | 1 | 1 | 1 | 1 |
| **P1D** | 0 | 1 | 1 | 0 | 1 | 1 | 1 | 1 | 0 | 1 | 1 | 0 |
| **P1E** | 0 | 1 | 1 | 1 | 1 | 1 | 0 | 1 | 0 | 0 | 0 | 0 |
| **P1F** | 0 | 0 | 1 | 0 | 0 | 0 | 1 | 0 | 1 | 0 | 1 | 0 |
| **P1G** | 0 | 0 | 1 | 0 | 1 | 1 | 0 | 1 | 0 | 1 | 0 | 0 |
| **P1H** | 0 | 0 | 1 | 0 | 0 | 1 | 0 | 0 | 0 | 1 | 0 | 0 |

| Plate 2 | 01 | 02 | 03 | 04 | 05 | 06 | 07 | 08 | 09 | 10 | 11 | 12 |
|---|---|---|---|---|---|---|---|---|---|---|---|---|
| **P2A** | 0 | 0 | 0 | 1 | 0 | 0 | 1 | 1 | 0 | 0 | 1 | 1 |
| **P2B** | 0 | 0 | 1 | 0 | 0 | 1 | 0 | 1 | 0 | 0 | 0 | 0 |
| **P2C** | 0 | 1 | 0 | 0 | 0 | 0 | 0 | 0 | 0 | 0 | 0 | 0 |
| **P2D** | 1 | 0 | 1 | 0 | 0 | 1 | 0 | 0 | 1 | 0 | 0 | 0 |
| **P2E** | 1 | 1 | 0 | 0 | 0 | 0 | 1 | 0 | 0 | 1 | 0 | 0 |
| **P2F** | 0 | 1 | 0 | 0 | 0 | 0 | 0 | 1 | 0 | 0 | 0 | 1 |
| **P2G** | 1 | 1 | 1 | 0 | 0 | 1 | 0 | 0 | 1 | 1 | 1 | 0 |
| **P2H** | 1 | 0 | 0 | 1 | 0 | 0 | 0 | 0 | 0 | 1 | 1 | 0 |

**b**

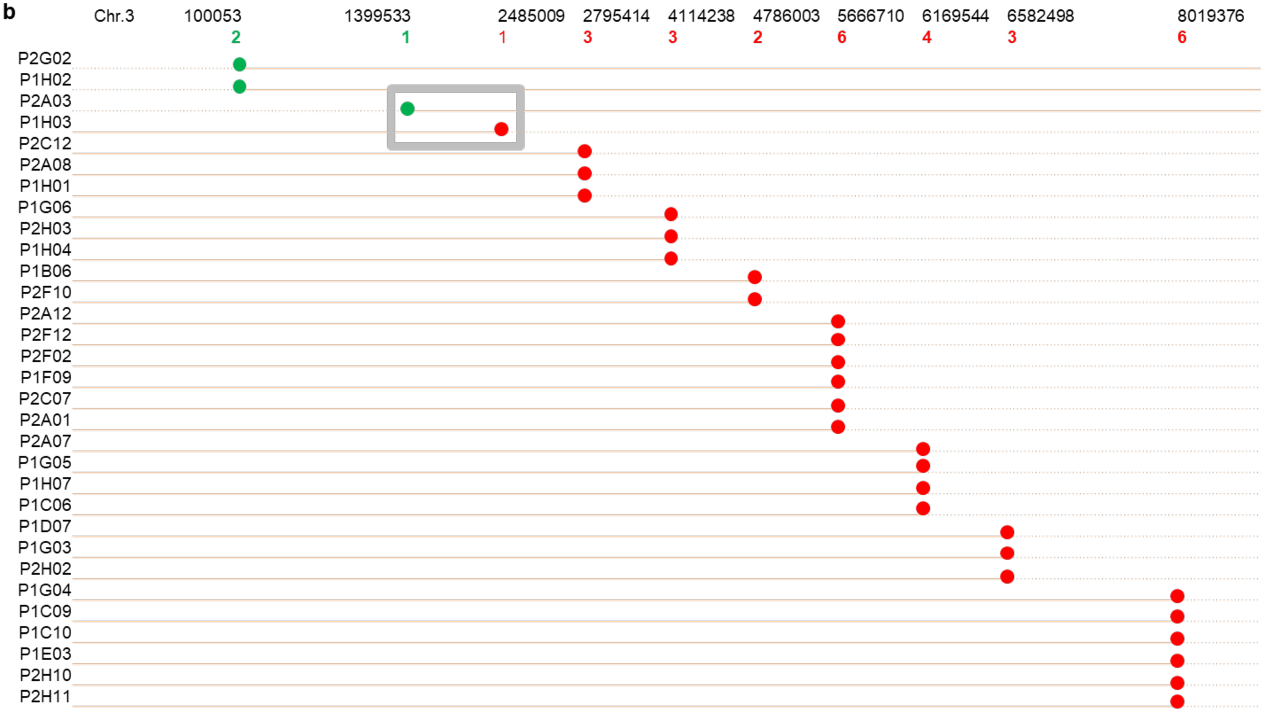


**Supplementary Figure 6. Definition of left and right boundaries of the QTL for RsE-sensitivity. a,** Binary phenotype layout for the Rad-seq library. The layout was designed according to a 96-well format, each well shows the value 1 (RsE-insensitive) or 0 (RsE-sensitive) of the RsE-induced ethylene response of individual $F_2$ plants of an ICE153 x ICE73 cross. Insensitive phenotypes are highlighted in grey. **b,** Diagram of 31 individual $F_2$ plants (code given on the left) containing informative recombination events with their genomic positions on chromosome 3 indicated on top. Green numbers indicate summarized frequency of recombination events occurring at the left boundary, red numbers indicate summarized frequency of recombination events occurring at the right boundary within the $F_2$ population. The grey box represents the rQTL-mapping region that is associated with RsE-induced ethylene responses.

| Line name | NASC stock number | Allele name | T-DNA position as verified by sequencing |
|---|---|---|---|
| SM_3_33092 | N119803 | *rlp32-2* | 12 bp downstream of start codon |
| SALK_137467C | N657024 | *rlp32-3* | 408 bp upstream of start codon |
| SM_3_33695 | N120406 | *rlp32-4* | 2 bp downstream of start codon |
| SM3_15851 | N106446 | *rlp32-5* | 487 bp downstream of start codon |

**Supplementary Figure 7. *Rlp32* mutant genotypes used in this study.** The position of the T-DNA or transposon insertion was verified in each line by flanking fragment sequencing. The mutant line FLAG_588C11 in the Ws-0 accession was published by Wang et al.[1] as *rlp32-1* and was not used in our studies.

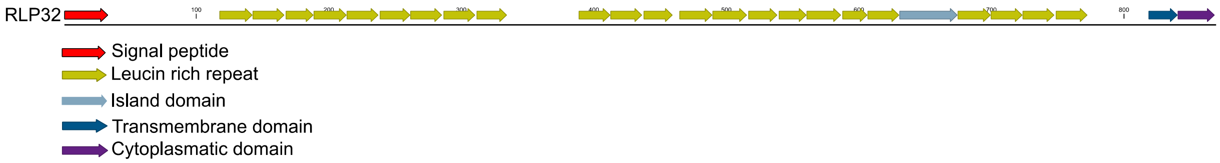


**Supplementary Figure 8. Schematic representation of the RLP32 protein structure.** RLP32 consists of a signal peptide, 23 LRR domains, an island domain, a transmembrane domain and a short cytoplasmatic tail. Protein domains identified by UniProt database are indicated in colors.

```
               10        20        30        40        50        60        70        80        90       100
Col-0  MKDSWNSTSIIPFTFSSLIFFLFTFDFQDVFGVPTKHLCRLEQRDALLELKKEFKIKKPCFDGLHPTTESWANNSDCCYWDGITCNDKSGEVLELDLSRS 100
Dog-4  ...XXXXX............................................................................................ 100
ICE21  .................................................................................................... 100
ICE73  ...............Y...XXX.....R......................................K.......................D......... 100

              110       120       130       140       150       160       170       180       190       200
Col-0  CLQSRFHSNSSLFTVLNLRFLTTLDLSYNYFSGQIPSCIENFSHLTTLDLSKNYFSGGIPSSIGNLSQLTFLDLSGNEFVGEMPFFGNMNQLTNLYVDSN 200
Dog-4  ...............Q.................................................................................... 200
ICE21  ...............Q.................................................................................... 200
ICE73  .............R.Q..P..................S.......................................D.........T............ 200

              210       220       230       240       250       260       270       280       290       300
Col-0  DLTGIFPLSLLNLKHLSDLSLSRNQFTGTLPSNMSSLSNLEYFEAWGNAFTGTLPSSLFTIASLTSINLRNNQLNGTLEFGNISSPSTLTVLDISNNNFI 300
Dog-4  .................................................................X.................................. 300
ICE21  .................................................................................................... 300
ICE73  E.............Q..............................................P..........H........................... 300

              310       320       330       340       350       360       370       380       390       400
Col-0  GPIPKSISKFINLQDLDLSHLNTQGPVDFSIFTNLKSLQLLNLSHLNTTTTIDLNALFSSHLNSIYSMDLSGNHVSATTKISVADHHPTQLISQLYLSGC 400
Dog-4  ..............................................................................I..................... 400
ICE21  .................................................................................................... 400
ICE73  ................................M...............................................S................... 400

              410       420       430       440       450       460       470       480       490       500
Col-0  GITEFPELLRSQHKMTNLDISNNKIKGQVPGWLWTLPKLIFVDLSNNIFTGFERSTEHGLSLITKPSMQYLVGSNNNFTGKIPSFICALRSLITLDLSDN 500
Dog-4  .................................................................................................... 500
ICE21  .................................................................................................... 500
ICE73  .................................................................................................... 500

              510       520       530       540       550       560       570       580       590       600
Col-0  NLNGSIPPCMGNLKSTLSFLNLRQNRLGGGLPRSIFKSLRSLDVGHNQLVGKLPRSFIRLSALEVLNVENNRINDTFPFWLSSLKKLQVLVLRSNAFHGP 600
Dog-4  ...................................................................................................A 600
ICE21  .................................................................................................... 600
ICE73  .................................................................................................... 600

              610       620       630       640       650       660       670       680       690       700
Col-0  IHHASFHTLRIINLSHNQFSGTLPANYFVNWNAMSSLMATEDRSQEKYMGDSFRYYHDSVVLMNKGLEMELVRILKIYTALDFSENKLEGEIPRSIGLLK 700
Dog-4  .................................................................................................... 700
ICE21  ............D....................................................................................... 700
ICE73  ............D....................................................................................... 700

              710       720       730       740       750       760       770       780       790       800
Col-0  ELHVLNLSSNAFTGHIPSSMGNLRELESLDVSQNKLSGEIPQELGNLSYLAYMNFSHNQLGGLVPGGTQFRRQNCSSFKDNPGLYGSSLEEVCLDIHAPA 800
Dog-4  .................................................................................................... 800
ICE21  ...............................................................E.................................... 800
ICE73  ..............................D..................................................................... 800

              810       820       830       840       850       860
Col-0  PQQHEPPELEEEDREVFSWIAAAIGFGPGIAFGLTIRYILVFYKPDWFMHTFGHLQPSAHEKRLRRKQ.                                869
Dog-4  ....................................G................................                                869
ICE21  ....................................G................................                                869
ICE73  ....................................G................................                                869
```

**Supplementary Figure 9. Alignment of RLP32 protein sequences.** Protein sequence alignment of RLP32 sequences from RsE-sensitive *Arabidopsis* accession Col-0 and RsE-insensitive accessions Dog-4, ICE21 and ICE73 highlighting accession-specific amino acid polymorphisms. Sequence data were retrieved from the 1001genomes website using the polymorph tool (http://polymorph.weigelworld.org/cgi-bin/webapp.cgi). X indicates sequence ambiguities in available data sets. Dots represent conserved amino acid residues.

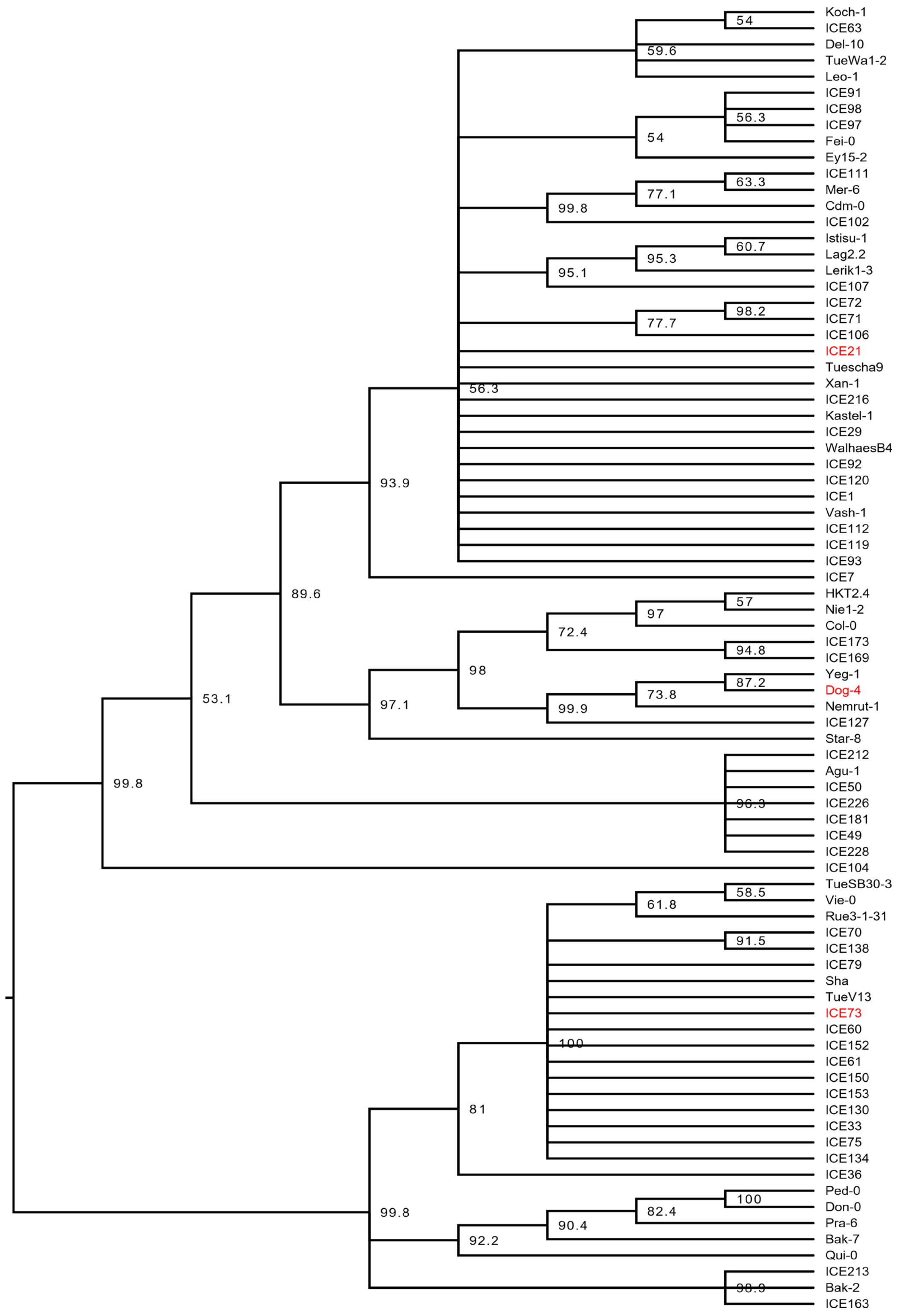


**Supplementary Figure 10. Phylogenetic tree of *Arabidopsis RLP32* gene sequences.** A phylogenetic tree of *RLP32* gene sequences from 80 *Arabidopsis* accessions was built using the neighbor-joining method with 1,000 bootstrap replications. The consensus tree was presented and bootstrap values over 50 % were indicated at the right side of each node. Insensitive ecotypes are given in red.

a

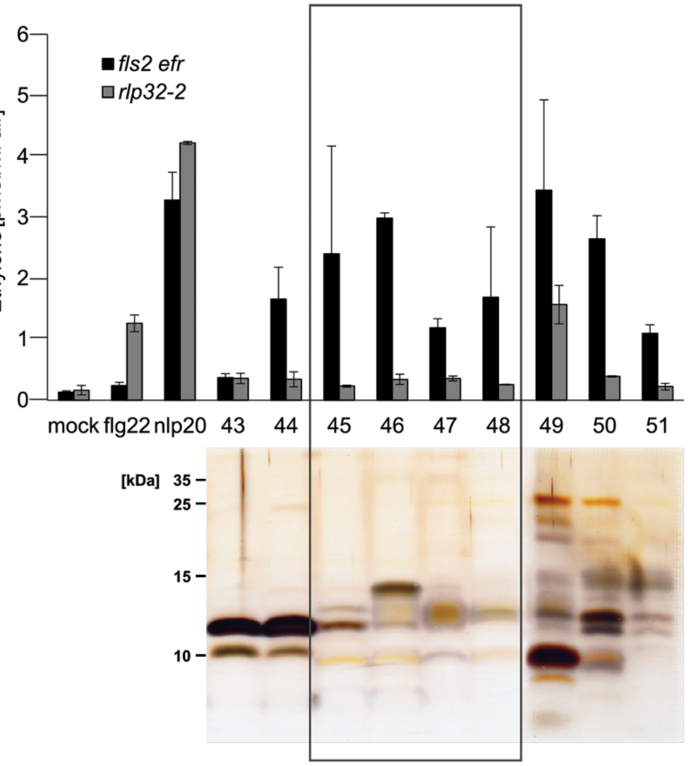


c

| | Fractions | | | | | |
|---|---|---|---|---|---|---|
| | 45 | 46 | 47 | 48 | | |
| Protein names | Number of peptides | | | | [kDa] | IP |
| Uncharacterized protein YibL | 15 | 23 | 25 | 17 | 13.70 | 9.49 |
| Uncharacterized protein YdcH | 17 | 18 | 16 | 14 | 8.88 | 9.3 |
| 50S ribosomal protein L24 | 16 | 13 | 14 | 10 | 11.32 | 10.21 |
| 50S ribosomal protein L22 | 10 | 10 | 13 | 17 | 12.23 | 10.23 |
| 50S ribosomal protein L13 | 13 | 13 | 11 | 12 | 16.02 | 9.91 |
| 50S ribosomal protein L17 | 10 | 8 | 12 | 11 | 14.36 | 11.05 |
| 50S ribosomal protein L6 | 9 | 8 | 13 | 11 | 18.90 | 9.71 |
| 50S ribosomal protein L30 | 9 | 12 | 12 | 6 | 6.54 | 10.96 |
| Spheroplast protein Y | 10 | 9 | 11 | 7 | 18.20 | 9.66 |
| 30S ribosomal protein S16 | 10 | 9 | 8 | 8 | 9.19 | 10.54 |
| Uncharacterized protein YciH | 9 | 8 | 7 | 10 | 11.40 | 9.38 |
| Chaperone protein Skp | 7 | 5 | 10 | 12 | 17.69 | 9.69 |
| Protein YqjC | 9 | 6 | 9 | 9 | 13.91 | 8.81 |
| 50S ribosomal protein L31 | 6 | 6 | 8 | 8 | 7.87 | 9.46 |
| Translation initiation factor IF1 | 6 | 7 | 7 | 7 | 8.25 | 9.22 |
| Acylphosphatase | 7 | 6 | 7 | 6 | 10.30 | 8.66 |
| Met repressor | 7 | 7 | 7 | 5 | 12.14 | 5.4 |
| 50S ribosomal protein L25 | 3 | 6 | 8 | 9 | 10.69 | 9.6 |
| 30S ribosomal protein S19 | 10 | 7 | 6 | 2 | 10.43 | 10.52 |
| 30S ribosomal protein S20 | 6 | 8 | 5 | 5 | 9.68 | 11.18 |

**Supplementary Figure 11. Identification of RLP32-dependent elicitor activity in fractionated *E. coli* proteins.** **a**, *E. coli* protein fractions separated by C8 reverse phase HPLC were assessed for eliciting ethylene production in *Arabidopsis fls2 efr* or *rlp32* plants. Treatment with water (mock), flg22 or nlp20 served as controls. **b**, Tricine-SDS-PAGE of proteins shown in (**a**). Proteins were visualized by silver staining. Boxed fractions 45-48 representing RLP32-dependent elicitor activity were analyzed by LC-MS/MS. **c**, List of proteins identified by LC-MS/MS. Shown are total numbers of peptides representing proteins identified by LC-MS/MS together with molecular masses (kDa) and isoelectric points (IP) of these proteins.

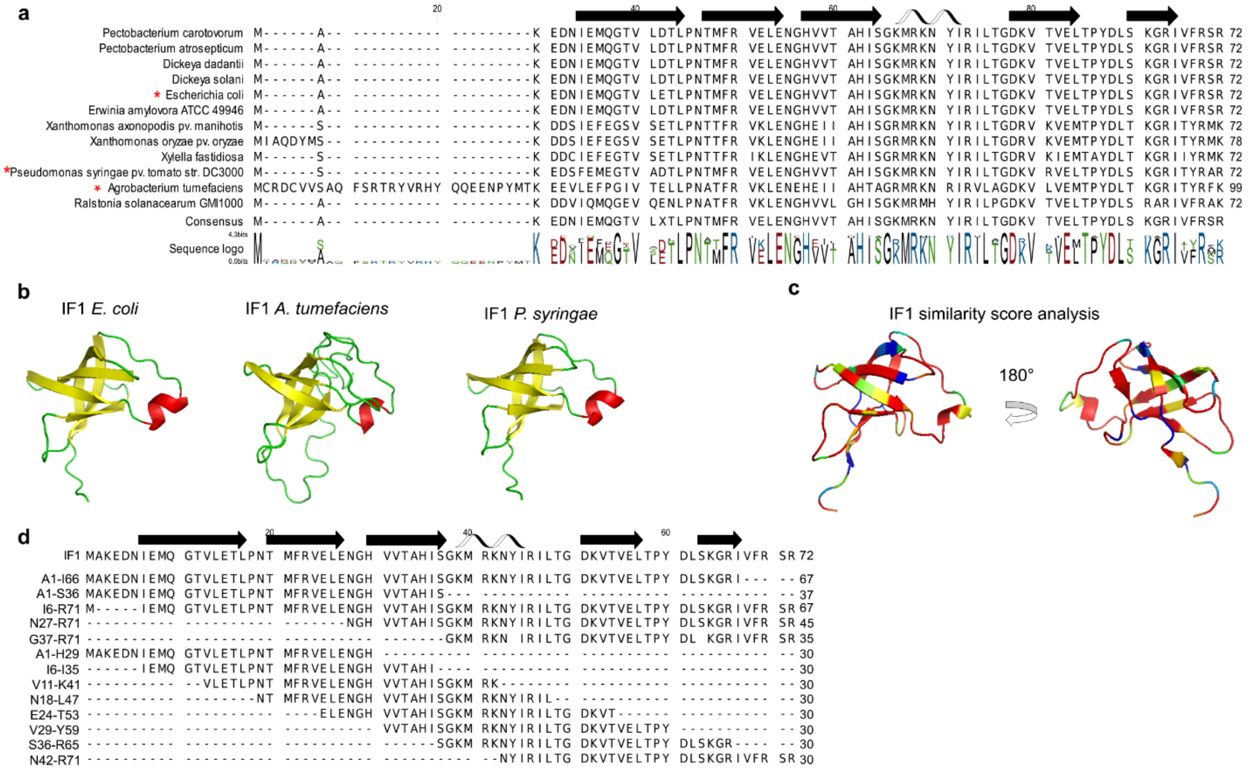


**Supplementary Figure 12. IF1 is conserved in *Proteobacteria*. a**, Alignment of IF1 amino acid sequences from different *Proteobacteria* species. Secondary structures (beta sheets indicated as black arrows and alpha helix indicated as grey helix) are indicated atop the protein sequences. Red asterisks indicate organisms from which IF1 was cloned and recombinantly expressed (see also Figure 2d). **b**, Iterative threading assembly refinement (I-TASSER) 3D structure prediction of IF1 derived from *E. coli*, *P. syringae* and *A. tumefaciens* as ribbon presentation (yellow indicates beta sheet, red indicates alpha helix). As a template, the NMR structure of IF1 derived from *E. coli* was used[2]. **c**, Ribbon representation of IF1, colored by conservation (blue indicates low B-factors; red indicates high B-factors). Conservation scores for each amino acid were calculated and mapped onto the IF1 structure with Easy Sequencing in PostScript (ESPript 2.2). Ribbon presentation was done with PYMOL. **d**, Alignment of amino acid sequences of *E. coli* IF1 deletion constructs and synthetic peptides used in Figure 2a and b with black arrows (beta sheets) and a helix symbol (alpha helix) indicating IF1 secondary structures.

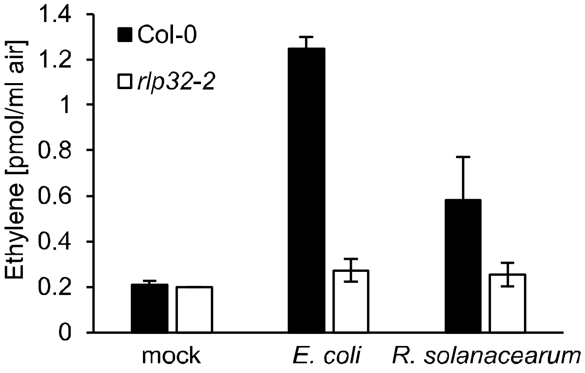


**Supplementary Figure 13: *R. solanacearum* IF1 displays RLP32-dependent activity.** Ethylene accumulation in *Arabidopsis* Col-0 and *rlp32-2* mutant plants after treatment with synthetic *E. coli* or *R. solanacearum* IF1, respectively. Water treatment served as control (mock). Bars represent means ± SD of three replicates.

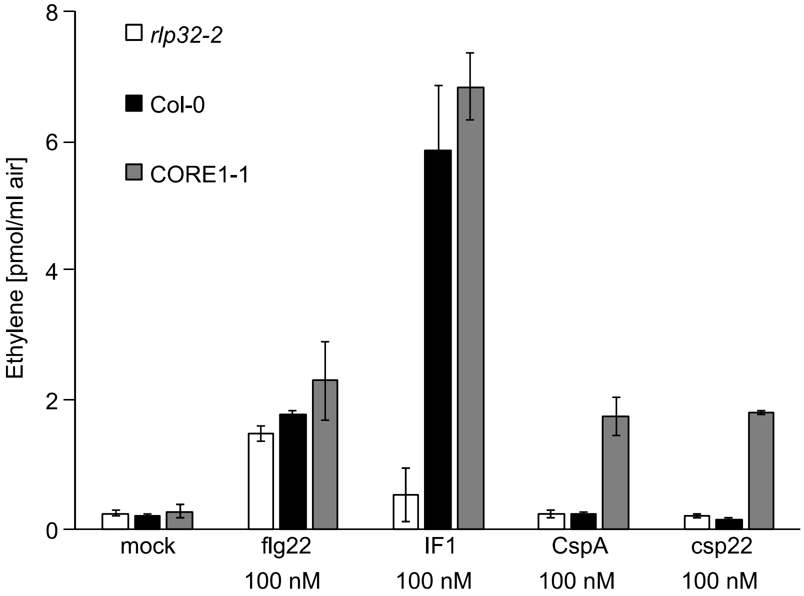


**Supplementary Figure 14. Bacterial cold shock protein CspA does not trigger ethylene production in *Arabidopsis*.** Col-0 wild-type plants, *rlp32* mutants or an *Arabidopsis* line stably expressing tomato cold shock receptor (CORE1-1)[3] were treated with the elicitors indicated and assessed for ethylene production. Water (mock) treatment served as control. Bars represent means ± SD of two replicates.

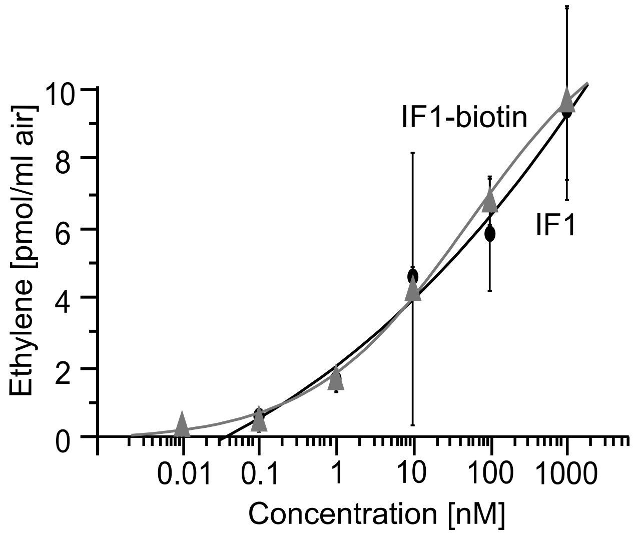


**Supplementary Figure 15. Biological activity of biotinylated IF1.** Ethylene accumulation in *Arabidopsis* Col-0 wild-type plants treated with IF1 recombinantly produced in *Pichia pastoris* (black) or treated with synthetic biotinylated IF1 (IF1-bio, grey). Data represent means ± SD of three replicates.

**Supplementary Table 1. Primers used for genotyping.**

| line | Primer name | Primer sequence (5' – 3') |
|---|---|---|
| *rlp32-2* (SM 3.3092) | L367 | AATTGTTCAAAACCGGTTGTG |
| | R1406 | CAGATTGAGTAGGGAAAGGGG |
| | Spm32 | GAATAAGAGCGTCCATTTTAGAGTG |
| *rlp32-3* (Salk_137467C) | L5 | CGGAATTGAAGACGTTCGTT |
| | R994 | TCACTGTTATTCGCCCATGA |
| | LBb1.3 | ATTTTGCCGATTTCGGAAC |
| *rlp32-4* (SM_3_33695) | L76 | AAATTGGGCTGATAAAATGGG |
| | R1186 | TCAATACAAGACGGGATTTGG |
| | Spm32 | GAATAAGAGCGTCCATTTTAGAGTG |
| *rlp32-5* (SM3_15851) | L528 | TGTTGACAATTCAACGCAGAG |
| | R1697 | AAATTTGGAAATGGATTTCGG |
| | Spm32 | GAATAAGAGCGTCCATTTTAGAGTG |

**Supplementary Table 2. Primers and synthetic genes used for cloning.**

| Template | Expression in | Primer name | Primer sequence (5' – 3') |
|---|---|---|---|
| RLP32 | *A. thaliana, N. benthamiana* | RLP32 endogenous promotor [-1597 bp] forward | GATTGCTTTGTGGAGTGGACTG |
| | | RLP32-ATG forward | ATGAAAGACTCTTGGAACTCAACGAG |
| | | RLP32 stop reverse | TTATTGCTTTCTCCTCAATCTTTTTTCATGTGC |
| | | RLP32 no stop reverse | TTGCTTTCTCCTCAATCTTTTTTCATGTGC |
| IF1, *E. coli* | | Start-forward | ATGGCCAAAGAAGACAATATTGAAATGCAAGG |
| | | Stop reverse | TCAGCGACTACGGAAGACAATGCGG |
| | | no stop reverse | GCGACTACGGAAGACAATGCGG |
| | | I6-R71 forward | ATGATTGAAATGCAAGGTACCGTTC |
| | | A1-I66 reverse | AATGCGGCCTTTGCTCAG |
| | | +Cys N-terminal forward | ATGTGCGCCAAAGAAGACAATATTG |
| IF1, *E. coli* | *Pichia pastoris* | EcoRI_IF1_fwd | AATTGAATTCATGGCCAAAGAAGACAATATTGAAATGCAAGG |
| | | NotI_IF1_nostop | AGAATTGCGGCCGCGCGACTACGGAAGACAATGCGG |
| | | IF1_K38E_for | ACTGCACACATCTCCGGTGAAATGCGCAAAAACTACATCC |
| | | IF1_K38E_rev | GATGTAGTTTTTGCGCATTTCACCGGAGATGTGTGCAGTAACC |
| | | IF1_R40E_for | GCACACATCTCCGGTAAAATGGAAAAAAACTACATCCGCATCCT |
| | | IF1_R40E_rev | AGGATGCGGATGTAGTTTTTTTCCATTTTACCGGAGATGTGTGC |
| | | IF1_R40L_for | CACACATCTCCGGTAAAATGCTCAAAAACTACATCCGCATCC |
| | | IF1_R40L_rev | AGGATGCGGATGTAGTTTTTGAGCATTTTACCGGAGATGTGTGC |
| | | IF1_K38R40K41L_for | TTACTGCACACATCTCCGGTCTAATGCTCTTAAACTACATCCGCATCCTG |
| | | IF1_K38R40K41L_rev | AGGATGCGGATGTAGTTTAAGAGCATTAGACCGGAGATGTGTGCAGT AACC |
| | | IF1_R40pK41_for | ATCTCCGGTAAAATGCGCCCGAAAAACTACATCCGCATCC |
| | | IF1_R40pK41_rev | ATGCGGATGTAGTTTTTCGGGCGCATTTTACCGGAGATG |
| IF1, *A. tumefaciens* C58 | | C58 forward | ATGTGCCGGGATTGTGTAG |
| | | C58 no stop reverse | CTTGAAGCGATAGGTGATGC |
| IF1, *P. syringae* | | DC3000 forward | ATGTCGAAAGAAGACAGCTTCGAAA |
| | | DC3000 no stop reverse | ACGAGCGCGGTAGGTGAT |
| CspA | *Pichia pastoris* | Synthetic gene construct | gaattcATGTCCGGTAAAATGACTGGTATCGTAAAATGGTTCAACGCTGA CAAAGGCTTCGGCTTCATCACTCCTGACGATGGCTCTAAAGATGTGTTC GTACACTTCTCTGCTATCCAGAACGATGGTTACAAATCTCTGGACGAAG GTCAGAAAGTGTCCTTCACCATCGAAAGCGGCGCTAAAGGCCCGGCAG CTGGTAACGTAACCAGCCTGTAAgcggccgc |
| CspA-IF1-Helix | *Pichia pastoris* | Synthetic gene construct | gaattcATGTCAGGGAAAATGACAGGAATCGTTAAGTGGTTCAATGCTG ACAAAGGCTTTGGCTTCATTACTCCAGATGATGGTAGTAAGGACGTAT TTGTGCATTTCTCTGCCATTCAATCCGGAAAGATGAGAAAGAACTACC TTGATGAAGGTCAGAAAGTCTCGTTTACCATAGAGTCTGGAGCTAAA GGTCCTGCAGCTGGTAACGTTACTAGCTTGgcggccgc |

### References


1 Wang, G. *et al.* A genome-wide functional investigation into the roles of receptor-like proteins in *Arabidopsis*. *Plant Physiol.* **147**, 503-517 (2008).

2 Sette, M. *et al.* The structure of the translational initiation factor IF1 from *E-coli* contains an oligomer-binding motif. *Embo J.* **16**, 1436-1443 (1997).

3 Wang, L. *et al.* The pattern-recognition receptor CORE of Solanaceae detects bacterial cold-shock protein. *Nat. Plants* **2**, 16185 (2016).
